# Discovering the mechanics of artificial and real meat

**DOI:** 10.1101/2023.06.04.543638

**Authors:** Skyler R. St. Pierre, Divya Rajasekharan, Ethan C. Darwin, Kevin Linka, Marc E. Levenston, Ellen Kuhl

## Abstract

Artificial meat is an eco-friendly alternative to real meat that is marketed to have a similar taste and feel. The mechanical properties of artificial meat significantly influence our perception of taste, but how precisely the mechanics of artificial meat compare to real meat remains insufficiently understood. Here we perform mechanical tension, compression, and shear tests on isotropic artificial meat (Tofurky® Plant-Based Deli Slices), anisotropic artificial meat (Daring™ Chick’n Pieces) and anisotropic real meat (chicken) and analyze the data using constitutive neural networks and automated model discovery. Our study shows that, when deformed by 10%, artificial and real chicken display similar maximum stresses of 21.0 kPa and 21.8 kPa in tension, -7.2 kPa and -16.4 kPa in compression, and 2.4 kPa and 0.9 kPa in shear, while the maximum stresses for tofurky were 28.5 kPa, -38.3 kP, and 5.5 kPa. To discover the mechanics that best explain these data, we consulted two constitutive neural networks of Ogden and Valanis-Landel type. Both networks robustly discover models and parameters to explain the complex nonlinear behavior of artificial and real meat for individual tension, compression, and shear tests, and for all three tests combined. When constrained to the classical neo Hooke, Blatz Ko, and Mooney Rivlin models, both networks discover shear moduli of 94.4 kPa for tofurky, 35.7 kPa for artificial chick’n, and 21.4 kPa for real chicken. Our results suggests that artificial chicken succeeds in reproducing the mechanical properties of real chicken across all loading modes, while tofurky does not, and is about three times stiffer. Strikingly, all three meat products display shear softening and their resistance to shear is about an order of magnitude lower than their resistance to tension and compression. We anticipate our study to inspire more quantitative, mechanistic comparisons of artificial and real meat. Our automated-model-discovery based approach has the potential to inform the design of more authentic meat substitutes with an improved perception of taste, with the ultimate goal to reduce environmental impact, improve animal welfare, and mitigate climate change, while still offering the familiar taste and texture of traditional meat.

Our source code, data, and examples will be available at https://github.com/LivingMatterLab/CANNs.

## 1 Motivation

Plant-based meat substitutes, or artificial meats, are an increasingly popular alternative to animal consumption. Driven by a growing push for environmental sustainability [5], public health, and animal welfare [37], the global market for plant-based meat substitutes is projected to reach $85 billion by 2030 from $4.6 billion in 2018 [43]. Nevertheless, there is persistant reluctance to adopt plant-based alternatives [33], in part due to the failure of meat replacements to adequately mimic the aesthetic and experiential qualities of conventional farmed meat [15, 34]. Indeed, approaches to engineered artificial meat products vary widely and result in a large array of appearances and textures [7, 17]. Robust characterization of the constitutive behavior of meat substitutes may well be the key to successfully engineer plant-based alternatives that approximate the properties of farmed meats [23].

The mechanical properties of artificial meat products influence our oral processing and inform our sensory perception [10, 36]. To establish quantitative insight into the mechanical behavior of plant-based meat substitutes, we compare two existing products, Tofurkey® Plant-Based Deli Slices and Daring™ Artificial Chick’n Pieces, to real chicken. We test all three meat products in tension, compression, and shear [4]. To provide an unbiased analysis, we then adopt constitutive neural networks [20, 26] to autonomously discover a material model that explains the relationship between stress and deformation [11, 27].

Constitutive neural networks are a new paradigm to discover the model, parameters, and experiments to best describe a material, entirely without human interaction [28]. In classical mechanics, the gold standard practice is to a priori select a model based on qualitative observations, for example, the shape of an experimental curve, the expected material behavior, or simply user preference [25]. However, the choice of the model inherently limits the fit to the data and, more importantly, the insight into the underlying physical phenomena that govern the material behavior. In standard machine learning, neural networks approximate the data to any degree of accuracy, but typically fail to provide any physical insight at all [1]. Constitutive neural networks combine the best of both words by hard-wiring constitutive constraints into the network design [14, 24, 29] and allow us to automatically screen a wide range of potential models while ensuring thermodynamic consistency [2, 27, 41].

To automatically discover the best model and parameters for artificial and real meat, we compared two distinct principal-stretch-based constitutive neural network architectures: the first is configured with strictly power law Ogden type terms [35] and the second is expanded to include exponential and logarithmic Valanis-Landel type terms [44, 45]. Furthermore, both architectures contain the popular neo Hooke [42], Blatz Ko [3], and Mooney Rivlin [32, 39] models as special cases. The Ogden type network allows us to extract the classical shear modulus for comparison with the literature [40], whereas the Valanis-Landel type network demonstrates to which extent additional exponential and logarithmic terms can improve model discovery [28]. We train both networks on new tension, compression, and shear data from two distinct meat substitutes and systematically compare them to real meat. This study is the first to characterize the mechanics of artificial meat products–fully autonomously and unbiased–without having to manually select a model and then fit its parameters to data.

## 2 Methods

### 2.1 Mechanical testing

We tested two types of artificial meat, Tofurkey® deli alices and Daring™ artificial chick’n, and, for comparison, one type of real meat, chicken. For each meat type, we tested *n* = 5 samples in tension, compression, and shear. Figure 1 illustrates our test setup for all nine mechanical tests.

**Figure 1:**
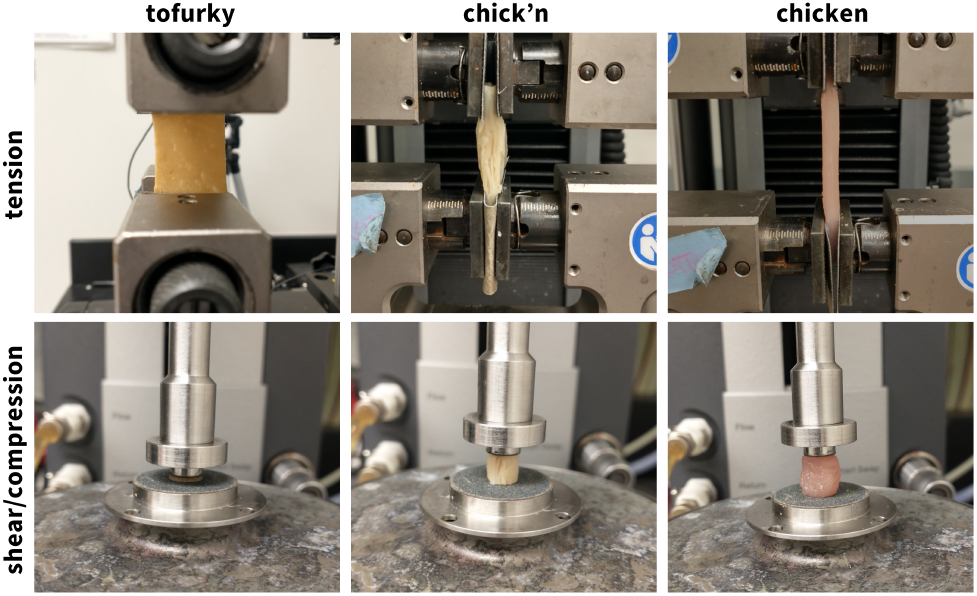
Mechanical testing of artificial and real meat. Tofurky® deli slices, Daring™ artificial chick’n, and real meat, from left to right, tested in uniaxial tension using the Instron 5848 test device, top, and in compression and shear using the AR-2000ex torsional rheometer, bottom. The artificial chick’n and real chicken were along the fiber direction in all testing modes.

#### 2.1.1 Sample Preparation

For the tension tests, we prepared strip samples of 2 cm in width to match the width of the specimen holder. We aligned the fiber direction of the artificial and real chicken with the direction of loading. For the compression and shear tests, we prepared cylindrical samples of 8 mm diameter using a biopsy punch to extract full-thickness cores from the center of each material. We aligned the fiber direction of the artificial and real chicken with the long axis of the cylindrical punch.

#### 2.1.2 Sample Testing

For all test modes, we tested the samples raw and at room temperature at 25^*◦*^C. We performed all uniaxial tension tests using an Instron 5848 (Instron, Canton, MA) with a 100N load cell, see Figure 1, top row. We mounted the sample, applied a small pre-load of 0.5 N, and calibrated the initial gage length *L*. We then increased the stretch quasi-statically at a rate of 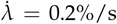 for *t* = 50 s to a total stretch of *λ* = 1.1 We performed all uniaxial compression and shear tests using an AR-2000ex torsional rheometer (TA Instruments, New Castle, DE), see Figure 1, bottom row. We mounted the sample, applied a small pre-load of 0.5 N, and calibrated the initial gage length *L*. We then compressed the tofurky samples quasi-statically at the rheometer’s minimum rate of 10 *µ*m/s and the real chicken and artificial chick’n samples at a rate of 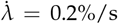 for *t* = 50 s, all to a total stretch of *λ* = 0.9. For the shear tests, we applied a small compressive pre-load and calibrated the initial gage length *L*. We then rotated the sample quasi-statically at 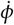at a shear rate of 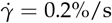 for *t* = 50 s, to a total shear of *γ* = 0.1. To prevent slippage of the samples during the shear tests, we used a sandpaper-covered base plate of 20 mm diameter and a sandpaper-covered top plate of 8 mm diameter.

#### 2.1.3 Analytical Methods and Data Processing

For each sample and each test mode, we used MATLAB (Mathworks, Natick, MA, USA) to smooth the curves using `smoothingspline` and `SmoothingParam = 1`. We interpolated the fit curve over 20 equally spaced points in the ranges 1.0 *≤ λ ≤* 1.1 for tension, 1.0 *≥ λ ≥* 0.9 for compression, and 0.0 *≤γ ≤*1.0 for shear. Finally, we averaged the five interpolated curves to obtain the mean, standard error of the mean, and standard error.

### 2.2 Kinematics

During testing, particles ***X*** of the undeformed sample map to particles ***x*** of the deformed sample via the deformation map ***φ*** such that ***x*** = ***φ***(***X***). Similarly, line elements of the dX of the undeformed sample map to line elements d***x*** of the deformed sample via the deformation gradient ***F*** such that d***x*** = ***F***·d***X***. The deformation gradient ***F*** is the gradient of the deformation map ***φ*** with respect to the undeformed coordinates ***X***. Its spectral representation introduces the principal stretches *λ*_*i*_ and the principal directions ***N***_*i*_ and ***n***_*i*_ in the undeformed and deformed configurations, where ***F*** · ***N***_*i*_ = *λ*_*i*_***n***_*i*_, and

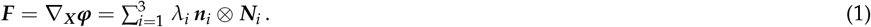

Here we assume that all samples are perfectly incompressible, and their Jacobian, *J* = det(***F***), always remains equal to one, *J* = 1. For simplicity, we also assume that all samples are isotropic and have three principal invariants,

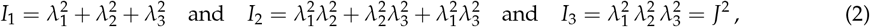

which are linear, quadratic, and cubic in terms of the principal stretches squared. The principal stretches depend on the type of experiment.

#### 2.2.1 Tension and compression

In the tension and compression experiments, we apply a stretch *λ* = *l*/*L*, that we calculate as the ratio between the current and initial sample lengths *l* and *L*. We can write the deformation gradient ***F*** in matrix representation as

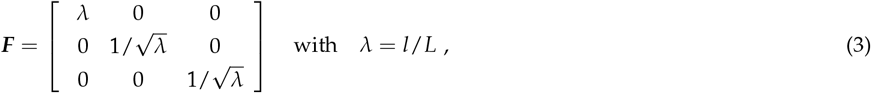

and immediately identify the principal stretches,

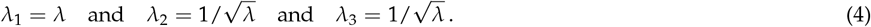

In tension and compression, the first and second invariants and their derivatives are

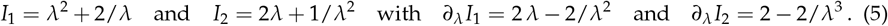

#### 2.2.2 Shear

In the shear experiment, we apply a torsion angle *ϕ*, that translates into the shear stress, *γ* = *r*/*L ϕ*, by multiplying it with the sample radius *r* and dividing by the initial sample length *L*. We can write the deformation gradient ***F*** in matrix representation as

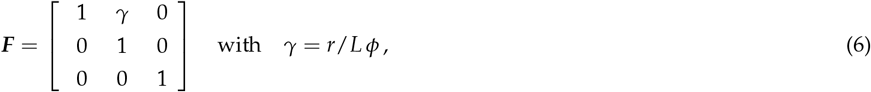

and calculate the principal stretches,

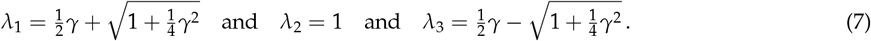

In shear, the first and second invariants and their derivatives are

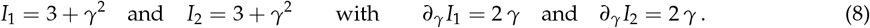

### 2.3 Constitutive Equations

Constitutive equations relate a stress like the Piola or nominal stress ***P***, the force per undeformed area that is commonly measured in experiments, to a deformation measure like the deformation gradient ***F***. For a hyperelastic material that satisfies the second law of thermodynamics, we can express the Piola stress, ***P*** = *∂ψ*(***F***)/*∂****F***, as the derivative of the Helmholtz free energy function *ψ*(***F***) with respect to the deformation gradient ***F***, modified by a pressure term, −*p **F***^**-t**^, to ensure perfect incompressibility,

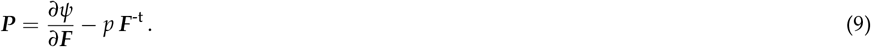

Here, the hydrostatic pressure, 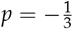 ***P:F*** acts as a Lagrange multiplier that that we determine from the boundary conditions. Instead of formulating the free energy function directly in terms of the deformation gradient *ψ*(***F***), we can either express it in terms of the invariants, *ψ*(*I*_1_, *I*_2_, *I*_3_), to yield the following expression for the Piola stress,

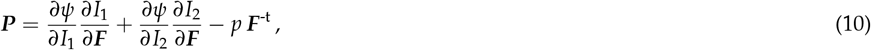

or in terms of the principal stretches, *ψ*(*λ*_1_, *λ*_2_, *λ*_3_), to result in the following Piola stress,

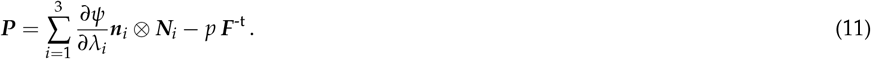

Here we focus on principal-stretch based constitutive models.

## 3 Neural network modeling

Motivated by these kinematic and constitutive considerations, we reverse-engineer two families of principal-stretch based neural networks that satisfy the conditions of thermodynamic consistency, material objectivity, material symmetry, incompressibility, constitutive restrictions, and polyconvexity by design [20, 41]. Instead of building these constraints into the loss function [8, 38], we hardwire them directly into our network input, output, architecture, and activation functions [27] to satisfy the fundamental laws of physics. We compare two different network architectures, an Ogden type network in Section 3.1 and a Valanis-Landel type network in Section 3.2. Special members of this family represent well-known constitutive models, including the neo Hooke [42], Blatz Ko [3], and Mooney Rivlin [32, 39] models, for which the network weights gain a clear physical interpretation [28, 40].

### 3.1 Ogden type neural network

Our first neural network is inspired by the Ogden model [35] and uses a free energy function that is parameterized in terms of the principal stretches, *λ*_1_, *λ*_2_, *λ*_3_,

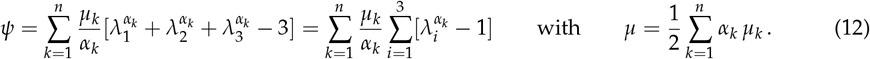

In this general form, the Ogden model consists of *n* terms and introduces 2*n* parameters, *n* stiffness-like parameters *µ*_*i*_ and *n* nonlinearity parameters *α*_*i*_, which collectively translate into the classical shear modulus *µ* from linear theory [19]. Motivated by the free energy function (12), we reverse-engineer a hyperelastic, perfectly incompressible, isotropic, principal-stretch-based Ogden type neural network [40]. The network takes the deformation gradient ***F*** for tension and compression (3) or shear (6) as input and computes the associated principal stretches *λ*_1_, *λ*_2_, *λ*_3_ using (4) or (7). From these stretches, it determines *n* = 20 Ogden terms, with fixed exponential coefficients, here ranging from *α*_1_ = *−*30 to *α*_*n*_ = +30 in increments of three, that make up the *n* nodes of the hidden layer of the model. The sum of all Ogden terms defines the strain energy function *ψ* as the network output. Figure 2 illustrates our Ogden type network with *n* = 20 nodes, for which the free energy function takes the following form,

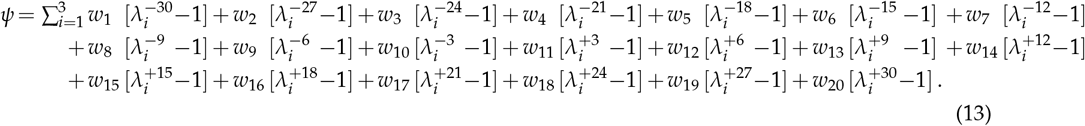

**Figure 2:**
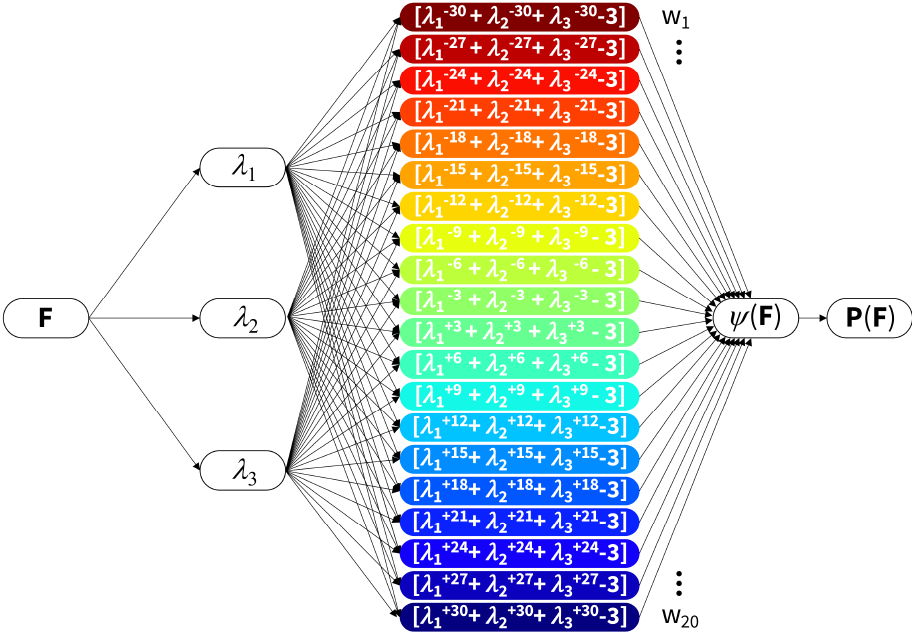
Ogden type constitutive neural network. The network represents a 20-term Ogden model with fixed exponents *α*_*k*_ ranging from -30 to +30 in increments of three. It takes the deformation gradient **F** as input and computes the three principal stretches *λ*_1_, *λ*_2_, *λ*_3_. From them, it calculates the Ogden terms for the 20 nodes of the hidden layer, multiplies them by the network weights *w*_*k*_, and sums up all terms to the strain energy function *ψ*(**F**) as output. The derivative of the strain energy function defines the Piola stress, **P** = *∂ψ*/*∂***F**, whose components *P*_11_ or *P*_12_ enter the normal force *N* or torque *M* in the loss function to minimize the error between the model and the tension, compression, and shear data.

Here, *w*_*k*_ are the network weights and 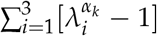 are the activation functions. From the free energy (13), we calculate the stress using equation (11), 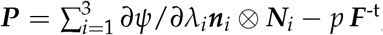, in terms of the derivative of the free energy *ψ* with respect to the stretches *λ*_*i*_,

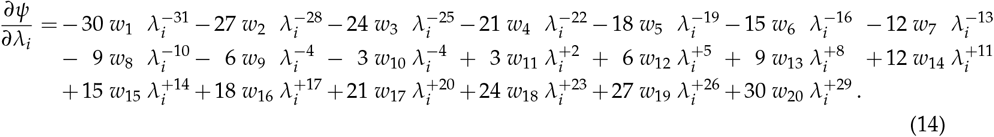

The network weights *w*_*k*_ are non-negative [2], and relate to the stiffness-like parameters *µ*_*k*_ and fixed exponential coefficients *α*_*k*_ as *w*_*k*_ = *µ*_*k*_/*α*_*k*_*≥*0. For our specific network with *n* = 20, the *α*_*k*_ coefficients are *α*_*k*_ = 3*k −n −*13 for *k≤* 10 and *α*_*k*_ = 3*k −n−* 10 for *k ≥*11. For this particular network, we recover the classical shear modulus *µ* from the linear theory in terms of the stiffness-like parameters *µ*_*k*_ as 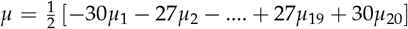 or, equivalently, in terms of the network weights *w*_*k*_ as 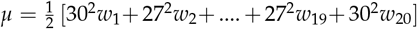 During training, our network autonomously identifies the best subset of activation functions from (2^*n*^ *−* 1) = 1, 048, 575 possible combinations of terms, and discovers the best model from more than a million possible models. At the same time, it naturally trains the weights of the less important terms to zero.

### 3.2 Valanis-Landel type neural network

Our second neural network is inspired by the Valanis Landel model [44], which postulates that the free energy function *ψ* can be expressed as the sum of any function subfunction *f* of the principal stretches, *λ*_1_, *λ*_2_, *λ*_3_,

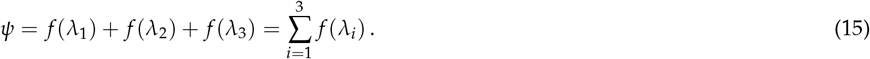

Comparisons with experimental data showed that logarithmic subfunctions *f* (*λ*_*i*_) = 2*µ* ln(*λ*_*i*_ *−*1) with derivatives *∂ f* /*∂λ*_*i*_ = 2*µ* ln(*λ*_*i*_) perform well in fitting uniaxial and biaxial data from natural rubber [44]. Other possible choices for the subfunction are the classical Ogden model [35] from equation (12), 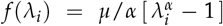 or exponential functions in the stretch, 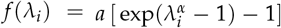, Motivated by the free energy function (15) and these considerations for the subfunction *f* (*λ*_*i*_), we reverse-engineer a hyperelastic, perfectly incompressible, isotropic, principal-stretch based Valanis-Landel type neural network. Similar to the Ogden type network, the Valanis-Landel type network takes the deformation gradient ***F*** for tension and compression (3) or shear (6) as input and computes the associated principal stretches *λ*_1_, *λ*_2_, *λ*_3_ using (4) or (7). From these stretches, it calculates the positive and negative second and fourth powers, (*◦*)^+2^, (*◦*)^+4^, (*◦*)^*−*2^, (*◦*)^+4^ and, for all four, calculates the Ogden type, exponential, and logarithmic terms. To give the network the additional freedom to discover larger Ogden type powers, we also include two terms, 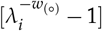 and 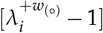 with trainable exponents, *−w*_(*◦*)_ and +*w*_(*◦*)_, resulting in a total of *n* = 4 × 3 + 2 = 14 terms. The sum of all Valanis-Landel terms defines the strain energy function *ψ* as the network output. Figure 3 illustrates our Valanis-Landel type network for which the free energy function takes the following form,

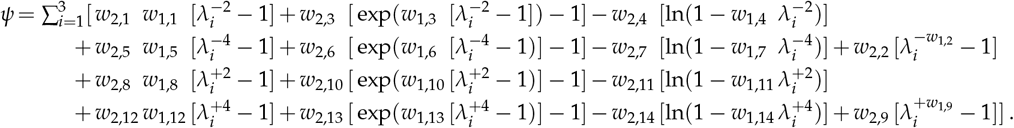

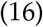

**Figure 3:**
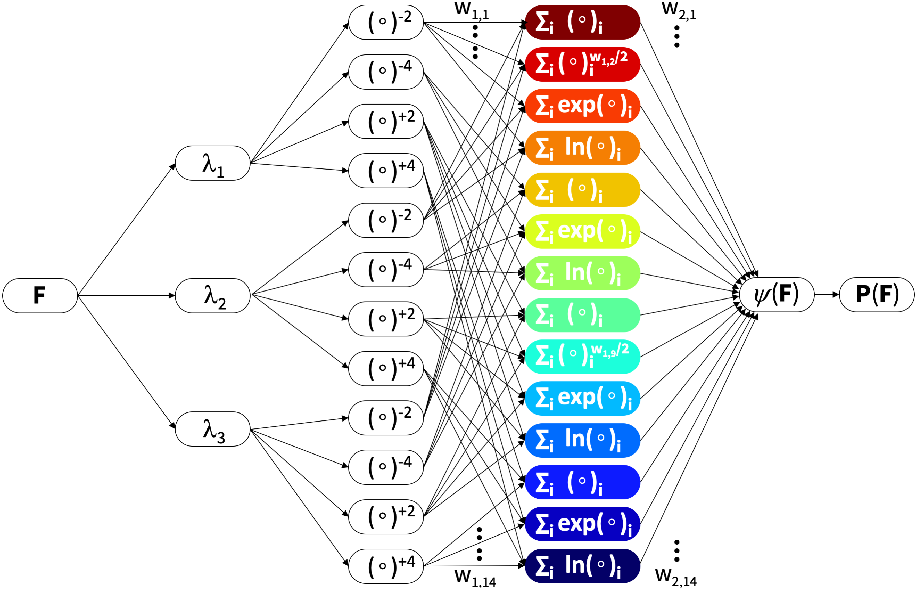
Valanis-Landel type neural network. The network takes the deformation gradient **F** as input and computes the three principal stretches *λ*_1_, *λ*_2_, *λ*_3_. From them, it calculates the positive and negative second and fourth powers, (*◦*)^+2^, (*◦*)^+4^, (*◦*)^*−*2^, (*◦*)^+4^ in the first hidden layer, multiplies them with their weights *w*_1,(*◦*)_, calculates their Ogden type terms, exponentials, logarithms in the second hidden layer, multiplies them with their weights *w*_2,(*◦*)_, and adds all terms to the strain energy function *ψ*(**F**) as output. The derivative of the strain energy function defines the Piola stress, **P** = *∂ψ*/*∂***F**, whose components *P*_11_ or *P*_12_ enter the normal force *N* or torque *M* in the loss function to minimize the error between the model and the tension, compression, and shear data.

Here *w*_1,*k*_ are the unit-less network weights from the first hidden layer and *w*_2,*k*_ are the stiffness-type network weights from the second hidden layer as shown in Figure 3. From the free energy (16), we calculate the stress using equation (11), 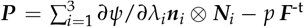 in terms of the derivative of the free energy *ψ* with respect to the principal stretches *λ*_*i*_,

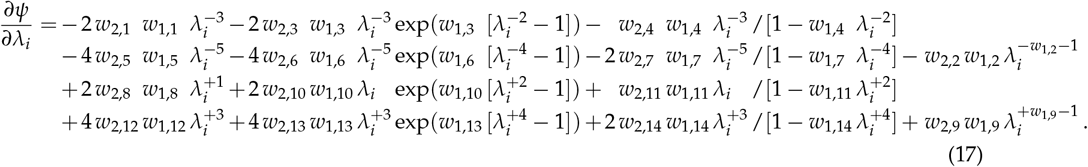

The network has 14 unit-less weights *w*_1,(*◦*)_ between the two hidden layers and 14 stiffness-type weights *w*_2,(*◦*)_ after the second hidden layer, with four redundant weights (*w*_2,1_*w*_1,1_), (*w*_2,5_*w*_1,5_), (*w*_2,8_*w*_1,8_), (*w*_2,12_*w*_1,12_), resulting in a total of 24 independent weights.

#### 3.2.1 Special cases

Our constitutive neural network in Figure 3 is a generalization of popular constitutive models. Specifically, we obtain the following one- and two-term models by setting the remaining network weights to zero.

The *neo Hooke* model [42] only uses the fixed exponent +2. Its free energy function is

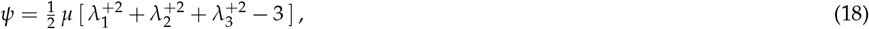

and its shear modulus is *µ* = 2 *w*_1,8_*w*_2,8_.

The *Blatz Ko* model [3] only uses the fixed exponent *−*2. Its free energy function is

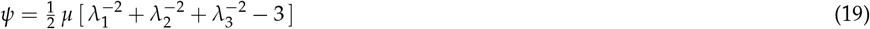

and its shear modulus is *µ* = 2*w*_1,1_*w*_2,1_.

The *Mooney Rivlin* model [32, 39] is a combination of both with fixed exponents *−*2 and +2. Its free energy function is

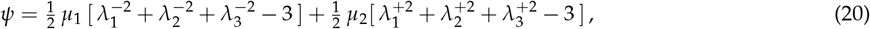

and its shear modulus is *µ* = *µ*_1_ + *µ*_2_ with *µ*_1_ = 2 *w*_1,1_*w*_2,1_ and *µ*_2_ = 2 *w*_1,8_*w*_2,8_.

The *general two term Ogden model* [35] uses two free exponents, *−α*_1_ = *−w*_1,2_ and +*α*_2_ = +*w*_1,9_. Its free energy function is

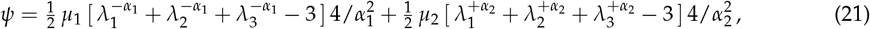

and its shear modulus is *μ* = *μ*_1_ + *μ*_2_ with 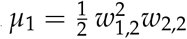 and 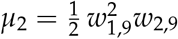

### 3.3 Loss function

Our constitutive neural networks learn the network weights, ***w*** = *w*_1_, …, *w*_*k*_, by minimizing a loss function *L* that penalizes the mean squared error, the *L*_2_-norm of the difference between model and data divided by the number of training points *n*_train_,

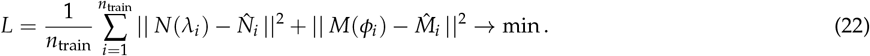

Specifically, the loss function contains the mean squared error between the normal forces of the model *N*(*λ*_*i*_) and the stretch-force pairs, 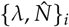 of the tension and compression experiments, and between the torque of the model *M*(*ϕ*_*i*_) and the torsion angle-torque pairs, 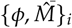 of the shear experiment. To reduce potential overfitting, we also study the effects of Ridge or L2 regularization,

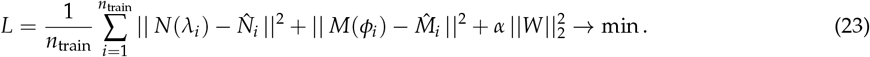

where *α* is the penalty parameter or regularization coefficient and 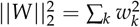 is the weighted L2 norm.

#### 3.3.1 Tension and compression

In the tension and compression experiments, the data are the recorded as stretch-force pairs, 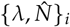 and the model output is the normal force as a function of the stretch *N*(*λ*_*i*_), which we calculate by integrating the normal component of the network model stress *P*_11_ across the cross section, d*A* = *r* d*r* d*θ*, of the sample [16],

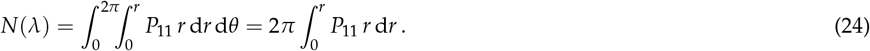

The normal stress *P*_11_ follows from the general stress equation (9), evaluated in the principal stretch space, *P*_*ii*_ = [*∂ψ*/*∂I*_1_][*∂I*_1_/*∂λ*_*i*_] + [*∂ψ*/*∂I*_2_][*∂I*_2_/*∂λ*_*i*_] − [1/*λ*_*i*_] *p*, for *i* = 1, 2, 3, using an invariant-based formulation. Here, *p* denotes the hydrostatic pressure that we determine from the zero normal stress condition, *P*_22_ = 0 and *P*_33_ = 0, as *p* = [2/*λ*][*∂ψ*/*∂I*_1_] + [2*λ* + 2/*λ*^2^][*∂ψ*/*∂I*_2_], to obtain the following explicit uniaxial stress-stretch relation for a hyperelastic, isotropic, incompressible material,

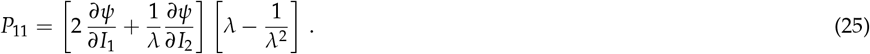

Alternatively, we can evaluate the general stress equation (9), in the principal stretch space, *P*_*ii*_ =*∂ψ*/*∂λ*_*i*_*− p*/*λ*_*i*_ for *i* = 1, 2, 3, using a principal-stretch-based formulation. Here, *p* denotes the hydrostatic pressure that we determine from the zero normal stress condition, *P*_22_ = 0 and *P*_33_ = 0, as *p* = *−*2*λ ∂ψ*/*∂λ*, to obtain an explicit equation for the normal stress,

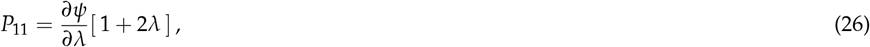

where *∂ψ*/*∂λ* is the derivative of the free energy *ψ* with respect to the applied stretch *λ* according to equations or (16) or (17). From equations (25) and (26) we conclude that the normal stress is constant across the cross section, independent of the radius *r*, and we can evaluate the normal force explicitly as,

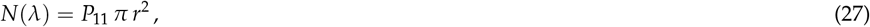

where the cross section area is *A* = *π r*^2^ for cylindrical compression samples and we replace this term by *A* = *b t* for rectangular tension samples. This implies that we can translate the recorded stretch-force pairs 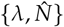 into stretch-stress pairs 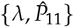 pairs with

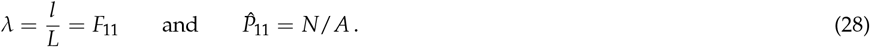

#### 3.3.2 Shear

In the shear experiment, the data are the recorded as torsion angle-torque pairs, 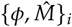 and the model output is the torque as a function of the torsion angle *M*(*ϕ*_*i*_), which we calculate by integrating the shear component of the network model stress *P*_12_ times its moment arm *r* across the cross section, d*A* = *r* d*r* d*θ*, of the sample [16],

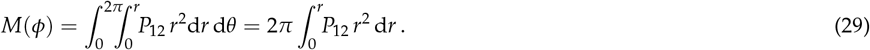

The shear stress *P*_12_ follows from the general stress equation (9), and we obtain the following explicit shear stress-stretch relation for a hyperelastic, isotropic, incompressible material,

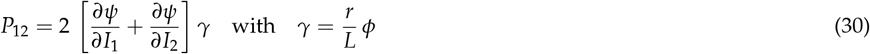

Unlike the normal stress in equations (25) and (26), the shear stress in equation (30) is not constant across the cross section but varies with the radius *r*. In other words, the derivatives d*ψ*(*r*)/d*I*_1_ and d*ψ*(*r*)/d*I*_2_ can be functions of the radius *r* through the explicit dependence on the shear strain, *γ* = *rφ*/*L*. This implies that we can obtain a relation between the torque *M* and the torsion angle *φ*,

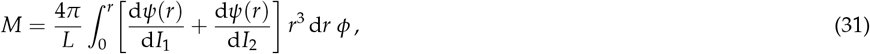

but we cannot simplify the integral any further. Here, we evaluate the integral along the radius *r* using numerical integration [12]. For example, using the trapezoidal rule, 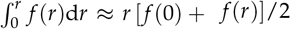, we obtain the following explicit relations between the torque *M*, torsion angle *ϕ*, shear *F*_12_, and shear stress *P*_12_,

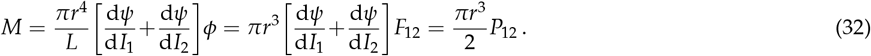

These equations are exact for the popular examples of the neo Hooke and Blatz Ko models with *ψ* = *µ* [*I*_1_ *−* 3] / 2 and *ψ* = *µ* [*I*_2_ *−* 3] / 2, for which we recover the classical explicit linear relations between the torque *M*, torsion angle *φ*, shear *γ*, and shear stress *P*_12_,

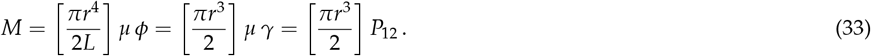

For this special case, we can translate the recorded torsion angle-torque pairs 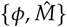 into shear stretch-stress pairs 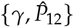 with

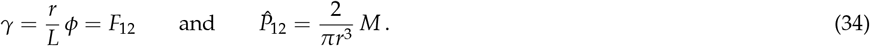

Traditionally, rheometer tests have been performed on stiff materials with small deformations, well in the linear regime, and have assumed this linear constitutive behavior a priori. For hyperelastic soft materials, with finite deformations and a nonlinear constitutive behavior [18], depending on whether the shear stress-stretch curve is concave or convex, equations (32) to (34) might very well under-or over-estimate the shear stress 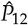

Motivated by these considerations, we reparameterize the loss function in terms of the mean squared error between the normal stresses of the model *P*_11_(*λ*_*i*_) and the stretch-stress pairs, 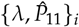 of the tension and compression experiments, and between the shear stress of the model *P*_12_(*γ*_*i*_) and the shear-stress pairs, 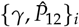 of the shear experiment, both scaled by the maximum absolute stress, max_*i*_ 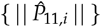 or max_*i*_ 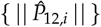 supplemented by L2 regularization,

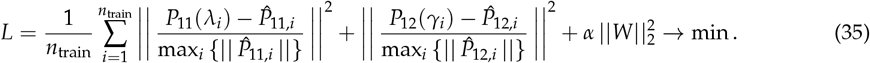

where *α* is the penalty parameter or regularization coefficient and 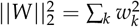is the weighted L2 norm. We train the network by minimizing the loss functions (35) or and learn the network parameters *w*_*k*_ using the ADAM optimizer, a robust adaptive algorithm for gradient-based first-order optimization, and constrain the weights to always remain non-negative, *w*_*k*_ *≥* 0.

### 3.4 Training and Testing Data

We train and test our Ogden and Valanis-Landel networks using tension, compression, and shear data from the tofurky, artificial chick’n, and real chicken as reported in Table 1. We perform single-mode training using a single loading case, either tension, compression, or shear, as training data and the remaining two cases as held-out test data. We perform multi-mode training using all three loading cases simultaneously as training data.

**Table 1:** Tofurky, artificial chick’n, and real chicken tested in tension, compression, and shear. Stresses are reported as means from the loading and unloading curves of *n* samples tested in the ranges 1.0 *≤ λ ≤* 1.1 for tension, 1.0 *≥ λ ≥* 0.9 for compression 0.0 *≤ γ ≤* 0.1 for shear.

| tofurky |  |  |  |  |  | chick'n |  |  |  |  |  | chicken |  |  |  |  |  |
| --- | --- | --- | --- | --- | --- | --- | --- | --- | --- | --- | --- | --- | --- | --- | --- | --- | --- |
| tension<br>*n* = 5 |  | comp<br>*n* = 5 |  | shear<br>*n* = 5 |  | tension<br>*n* = 5 |  | comp<br>*n* = 5 |  | shear<br>*n* = 5 |  | tension<br>*n* = 5 |  | comp<br>*n* = 5 |  | shear<br>*n* = 5 |  |
| $\lambda$<br>[-] | $P_{11}$<br>[kPa] | $\lambda$<br>[-] | $P_{11}$<br>[kPa] | $\gamma$<br>[-] | $P_{12}$<br>[kPa] | $\lambda$<br>[-] | $P_{11}$<br>[kPa] | $\lambda$<br>[-] | $P_{11}$<br>[kPa] | $\gamma$<br>[-] | $P_{12}$<br>[kPa] | $\lambda$<br>[-] | $P_{11}$<br>[kPa] | $\lambda$<br>[-] | $P_{11}$<br>[kPa] | $\gamma$<br>[-] | $P_{12}$<br>[kPa] |
| 1.000 | 0.000 | 1.000 | 0.000 | 0.000 | 0.000 | 1.000 | 0.000 | 1.000 | 0.000 | 0.000 | 0.000 | 1.000 | 0.000 | 1.000 | 0.000 | 0.000 | 0.000 |
| 1.005 | 2.498 | 0.995 | -1.602 | 0.005 | 0.525 | 1.005 | 1.826 | 0.995 | -0.616 | 0.005 | 0.306 | 1.005 | 1.579 | 0.995 | -1.190 | 0.005 | 0.127 |
| 1.010 | 4.536 | 0.990 | -3.230 | 0.010 | 1.019 | 1.010 | 3.303 | 0.990 | -1.216 | 0.010 | 0.552 | 1.010 | 2.802 | 0.990 | -2.316 | 0.010 | 0.215 |
| 1.015 | 6.358 | 0.985 | -4.830 | 0.015 | 1.436 | 1.015 | 4.614 | 0.985 | -1.746 | 0.015 | 0.755 | 1.015 | 3.834 | 0.985 | -3.259 | 0.015 | 0.281 |
| 1.020 | 8.062 | 0.980 | -6.428 | 0.020 | 1.819 | 1.020 | 5.853 | 0.980 | -2.246 | 0.020 | 0.931 | 1.020 | 4.772 | 0.980 | -4.207 | 0.020 | 0.345 |
| 1.025 | 9.706 | 0.975 | -8.074 | 0.025 | 2.182 | 1.025 | 7.041 | 0.975 | -2.713 | 0.025 | 1.095 | 1.025 | 5.691 | 0.975 | -5.221 | 0.025 | 0.401 |
| 1.030 | 11.242 | 0.970 | -9.717 | 0.030 | 2.502 | 1.030 | 8.171 | 0.970 | -3.077 | 0.030 | 1.241 | 1.030 | 6.593 | 0.970 | -6.118 | 0.030 | 0.451 |
| 1.035 | 12.752 | 0.965 | -11.405 | 0.035 | 2.823 | 1.035 | 9.243 | 0.965 | -3.433 | 0.035 | 1.377 | 1.035 | 7.513 | 0.965 | -7.007 | 0.035 | 0.497 |
| 1.040 | 14.208 | 0.960 | -13.121 | 0.040 | 3.114 | 1.040 | 10.265 | 0.960 | -3.845 | 0.040 | 1.503 | 1.040 | 8.454 | 0.960 | -7.931 | 0.040 | 0.538 |
| 1.045 | 15.620 | 0.955 | -14.854 | 0.045 | 3.383 | 1.045 | 11.288 | 0.955 | -4.246 | 0.045 | 1.616 | 1.045 | 9.408 | 0.955 | -8.725 | 0.045 | 0.577 |
| 1.050 | 16.990 | 0.950 | -16.668 | 0.050 | 3.640 | 1.050 | 12.275 | 0.950 | -4.631 | 0.050 | 1.722 | 1.050 | 10.402 | 0.950 | -9.439 | 0.050 | 0.616 |
| 1.055 | 18.313 | 0.945 | -18.538 | 0.055 | 3.879 | 1.055 | 13.237 | 0.945 | -5.011 | 0.055 | 1.821 | 1.055 | 11.424 | 0.945 | -10.196 | 0.055 | 0.648 |
| 1.060 | 19.573 | 0.940 | -20.434 | 0.060 | 4.105 | 1.060 | 14.208 | 0.940 | -5.335 | 0.060 | 1.913 | 1.060 | 12.450 | 0.940 | -10.958 | 0.060 | 0.679 |
| 1.065 | 20.797 | 0.935 | -22.422 | 0.065 | 4.312 | 1.065 | 15.169 | 0.935 | -5.613 | 0.065 | 1.995 | 1.065 | 13.516 | 0.935 | -11.694 | 0.065 | 0.713 |
| 1.070 | 22.036 | 0.930 | -24.485 | 0.070 | 4.511 | 1.070 | 16.111 | 0.930 | -5.907 | 0.070 | 2.066 | 1.070 | 14.618 | 0.930 | -12.413 | 0.070 | 0.743 |
| 1.075 | 23.209 | 0.925 | -26.594 | 0.075 | 4.697 | 1.075 | 17.001 | 0.925 | -6.205 | 0.075 | 2.130 | 1.075 | 15.737 | 0.925 | -13.122 | 0.075 | 0.770 |
| 1.080 | 24.342 | 0.920 | -28.808 | 0.080 | 4.870 | 1.080 | 17.869 | 0.920 | -6.475 | 0.080 | 2.191 | 1.080 | 16.913 | 0.920 | -13.823 | 0.080 | 0.798 |
| 1.085 | 25.427 | 0.915 | -31.048 | 0.085 | 5.038 | 1.085 | 18.696 | 0.915 | -6.708 | 0.085 | 2.246 | 1.085 | 18.080 | 0.915 | -14.430 | 0.085 | 0.822 |
| 1.090 | 26.513 | 0.910 | -33.375 | 0.090 | 5.190 | 1.090 | 19.504 | 0.910 | -6.908 | 0.090 | 2.298 | 1.090 | 19.273 | 0.910 | -15.015 | 0.090 | 0.844 |
| 1.095 | 27.542 | 0.905 | -35.823 | 0.095 | 5.337 | 1.095 | 20.285 | 0.905 | -7.074 | 0.095 | 2.351 | 1.095 | 20.547 | 0.905 | -15.710 | 0.095 | 0.869 |
| 1.100 | 28.543 | 0.900 | -38.297 | 0.100 | 5.465 | 1.100 | 20.999 | 0.900 | -7.231 | 0.100 | 2.395 | 1.100 | 21.803 | 0.900 | -16.446 | 0.100 | 0.888 |

## 4 Results

### 4.1 Experimental results

Figure 4 reports the tension, compression, and shear data for the tofurky, artificial chick’n, and real chicken converted into stretch-stress and shear-stress pairs using equations (28) and (34) with trapezoidal-rule type numerical integration. The means *±* standard error of the means reveal that there is a relatively small sample-to-sample variation. Notably, tofurky displays the stiffest response of all three materials, in all three modes, in tension, compression, and shear. Artificial chick’n and real chicken exhibit similar mechanical behaviors. We use the mean curves and values reported in Table 1 to train and test our neural networks.

**Figure 4:**
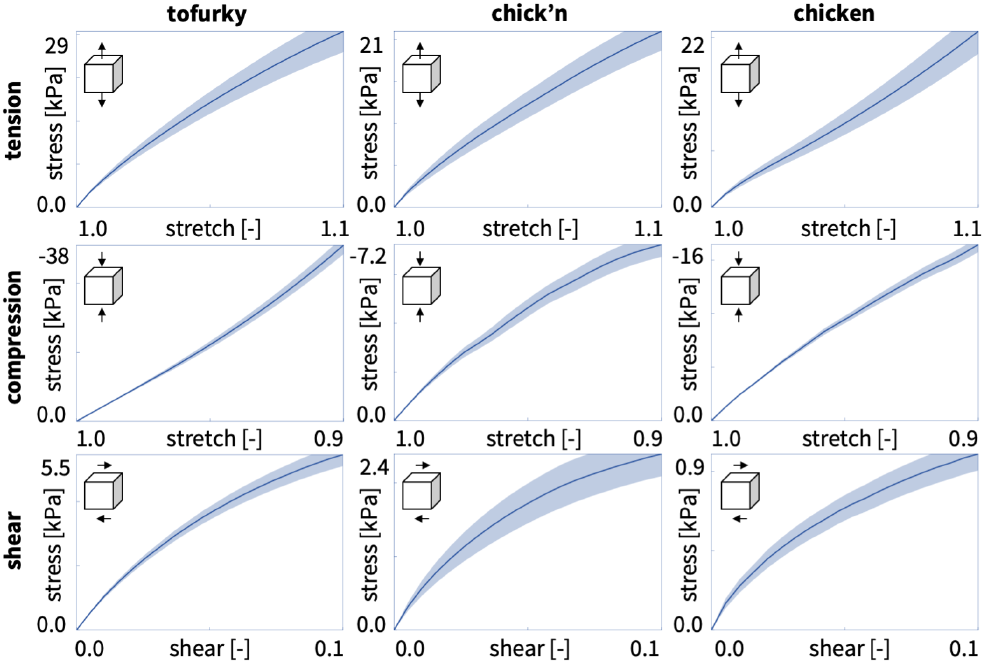
Tofurky, artificial chick’n, and real chicken tested in tension, compression, and shear. Stresses are reported as means *±* standard error of the means from the loading and unloading curves of *n* = 5 samples tested in the ranges 1.0 *≤ λ ≤* 1.1 for tension, 1.0 *≥ λ ≥* 0.9 for compression 0.0 *≤ γ ≤* 0.1 for shear.

### 4.2 Ogden type mechanics of artificial and real meat

Figure 5 illustrates the automatically discovered constitutive models for tofurky using the twenty term isotropic, perfectly incompressible Ogden type network from Figure 2. The three rows illustrate the Piola stress as a function of the stretch or shear for tension, compression, and shear. The first three columns indicate single-mode training for tension, compression, and shear, with training on the diagonal and testing on the off-diagonal. The last column shows the results of multi-mode training with all loading modes as training data. Each graph reports the goodness of fit 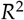 for either training or testing. The circles indicate the experimental data, while the color-coded regions designate the contributions of the twenty model terms to the free energy function *ψ*. Warm red-type colors indicate that the exponential power is negative, while cold blue-type colors indicate that the exponential power is positive. Table 2 shows the discovered weights and shear modulus of 80.08 kPa for multi-mode training. First, we observe that for single-mode training, the model succeeds in fitting the individual sets of training data with 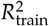values of 0.9940, 0.9997, and 0.9826 for tension, compression, and shear. Second, for single-mode training, the model is unable to predict all test data well. Third, the network is able to find an adequate fit of the data for multi-mode training, with training fit 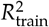 of 0.8627, 0.9641, and 0.8337 for tension, compression, and shear. Fourth, single-mode training with tension finds negative red-type terms, while the other modes find both negative and positive terms. Lastly, when trained with only the tension data, the network discovers a single negative term, the orange 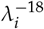 term in the left column, while for the other loading modes, it discovers a wide range and number of terms.

**Table 2:** Tofurky, artificial chick’n, and real chicken and discovered Ogden type models and parameters. Models and parameters are discovered for simultaneous training with tension, compression, and shear data using the Ogden type neural network from Figure 2. Summary of the weights *w*_*k*_, shear moduli 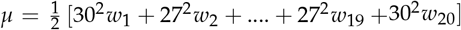 and goodness of fit *R*^2^ for training in tension, compression, and shear.

| | tofurky<br>ten+com+shr<br>$n = 5, 5, 5$ | chick'n<br>ten+com+shr<br>$n = 5, 5, 5$ | chicken<br>ten+com+shr<br>$n = 5, 5, 5$ |
| --- | --- | --- | --- |
|  | [kPa] | [kPa] | [kPa] |
| $w_1$ | 0.0080 | 0.0000 | 0.0141 |
| $w_2$ | 0.0003 | 0.0000 | 0.0000 |
| $w_3$ | 0.0003 | 0.0000 | 0.0000 |
| $w_4$ | 0.0146 | 0.0000 | 0.0000 |
| $w_5$ | 0.0358 | 0.0000 | 0.0000 |
| $w_6$ | 0.0575 | 0.0000 | 0.0000 |
| $w_7$ | 0.0784 | 0.0000 | 0.0000 |
| $w_8$ | 0.0962 | 0.0000 | 0.0000 |
| $w_9$ | 0.1060 | 0.0000 | 0.0000 |
| $w_{10}$ | 0.0834 | 0.0000 | 0.0000 |
| $w_{11}$ | 0.0920 | 0.0000 | 0.0000 |
| $w_{12}$ | 0.1280 | 0.0000 | 0.0000 |
| $w_{13}$ | 0.1282 | 0.0000 | 0.0000 |
| $w_{14}$ | 0.1166 | 0.0000 | 0.0000 |
| $w_{15}$ | 0.0972 | 0.0000 | 0.0000 |
| $w_{16}$ | 0.0714 | 0.0132 | 0.0000 |
| $w_{17}$ | 0.0402 | 0.0239 | 0.0000 |
| $w_{18}$ | 0.0044 | 0.0276 | 0.0000 |
| $w_{19}$ | 0.0001 | 0.0244 | 0.0000 |
| $w_{20}$ | 0.0000 | 0.0150 | 0.0334 |
| | $\mu = 80.08 \text{ kPa}$ | $\mu = 31.00 \text{ kPa}$ | $\mu = 21.39 \text{ kPa}$ |
| $R_t^2$ | 0.8627 | 0.6079 | 0.3226 |
| $R_c^2$ | 0.9641 | 0.7627 | 0.3424 |
| $R_s^2$ | 0.8337 | 0.8567 | 0.0000 |

**Figure 5:**
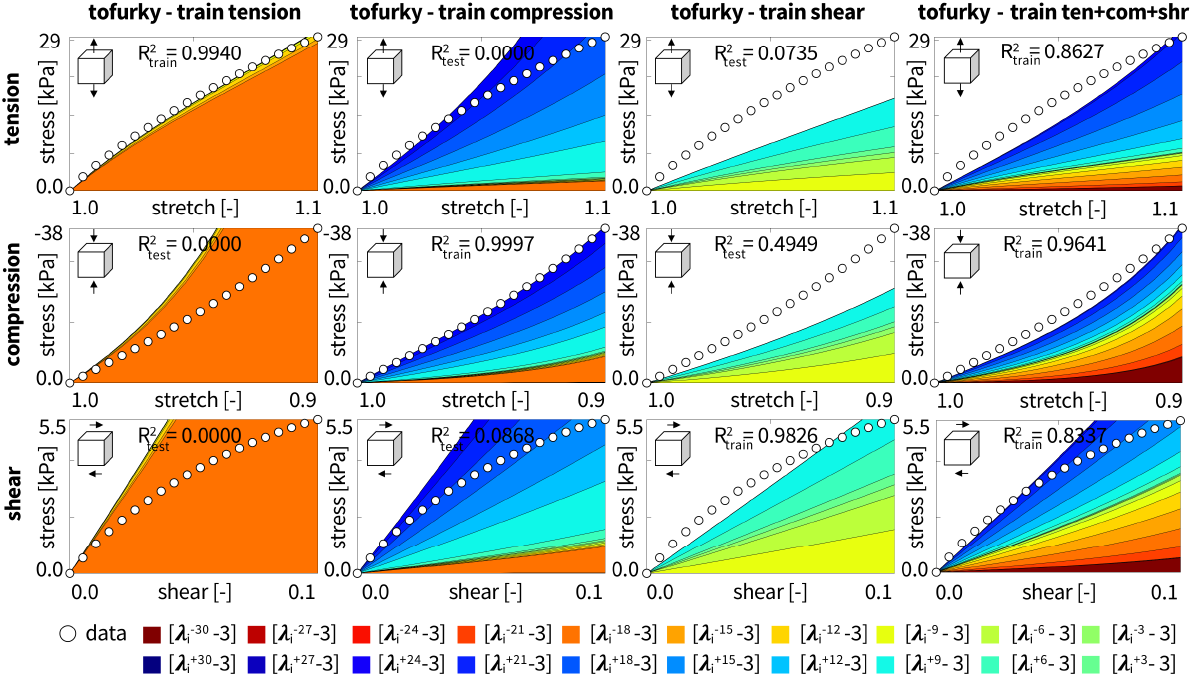
Tofurky data and Ogden type model. Nominal stress as a function of stretch or shear strain for the network with 20 nodes from Figure 2. Training individually with tension, compression, or shear data from the tofurky, as well as with all three load cases simultaneously. Circles represent the experimental data. Color-coded regions designate the contributions of the 20 model terms to the stress function according to Figure 2. Coefficients of determination *R*^2^ indicate goodness of fit for train and test data.

Figure 6 illustrates the automatically discovered models for artificial chick’n using the twenty term isotropic, perfectly incompressible Ogden type network from Figure 2. Table 2 shows the discovered weights and shear modulus of 31.00 kPa for multi-mode training. First, we note that for single-mode training, the model succeeds in fitting the individual sets of training data with values 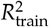 of 0.9967, 0.9409, and 0.9705 for tension, compression, and shear. Second, for single-mode training, the model performs well in predicting the shear data for training on compression with 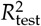 of 0.9682. Third, the network finds a moderately good fit of the data for multi-mode training, with training fit 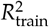of 0.6079, 0.7627, and 0.8567 for tension, compression, and shear. Fourth, single-mode training with tension or shear finds negative red-type terms, while single-mode training with compression or multi-mode training finds positive blue-type terms. Lastly, all types of training discover a small subset of the twenty total possible terms even without additional regularization, with between one to five terms visibly contributing to the stress function. Multimode training discovers only five positive terms from 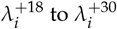

**Figure 6:**
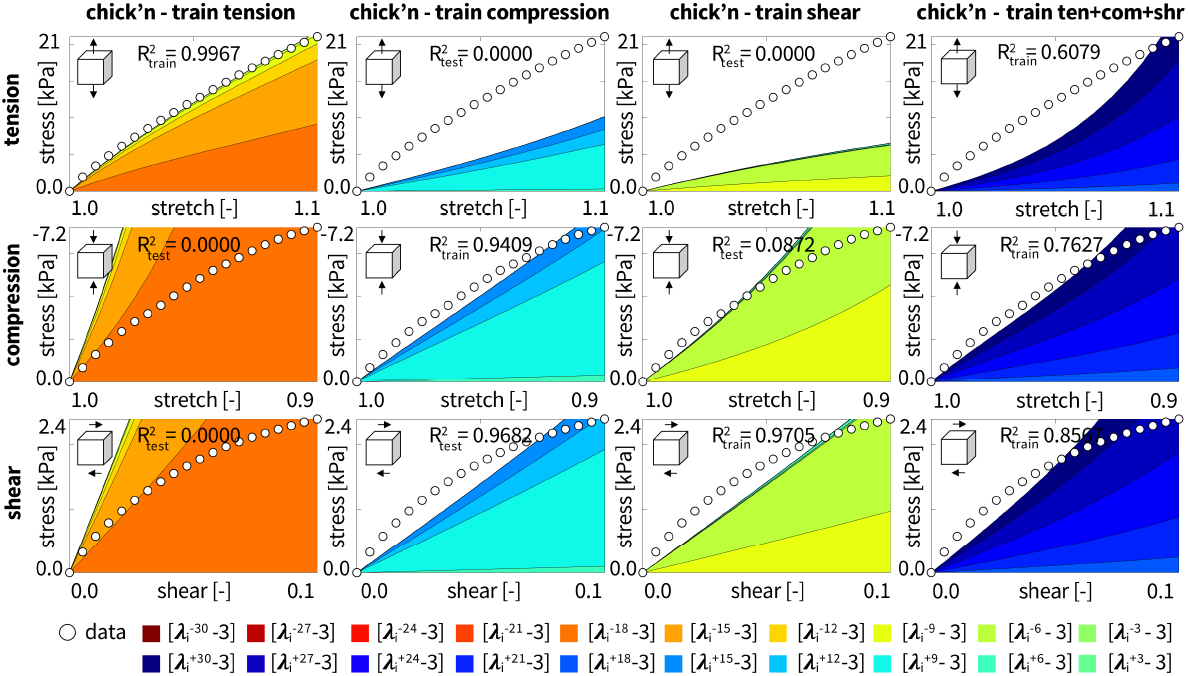
Artificial chick’n data and Ogden type model. Nominal stress as a function of stretch or shear strain for the network with 20 nodes from Figure 2. Training individually with tension, compression, or shear data from the artificial chick’n, as well as with all three load cases simultaneously. Circles represent the experimental data. Color-coded regions designate the contributions of the 20 model terms to the stress function according to Figure 2. Coefficients of determination *R*^2^ indicate goodness of fit for train and test data.

Figure 7 illustrates the automatically discovered models for real chicken using the twenty term isotropic, perfectly incompressible Ogden type network from Figure 2. Table 2 shows the discovered weights and shear modulus of 21.39 kPa for multi-mode training. First, we observe that for single-mode training, the principal-stretch-based model succeeds in fitting the individual sets of training data with 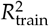values of 0.9994, 0.9764, and 0.9727 for tension, compression, and shear.

**Figure 7:**
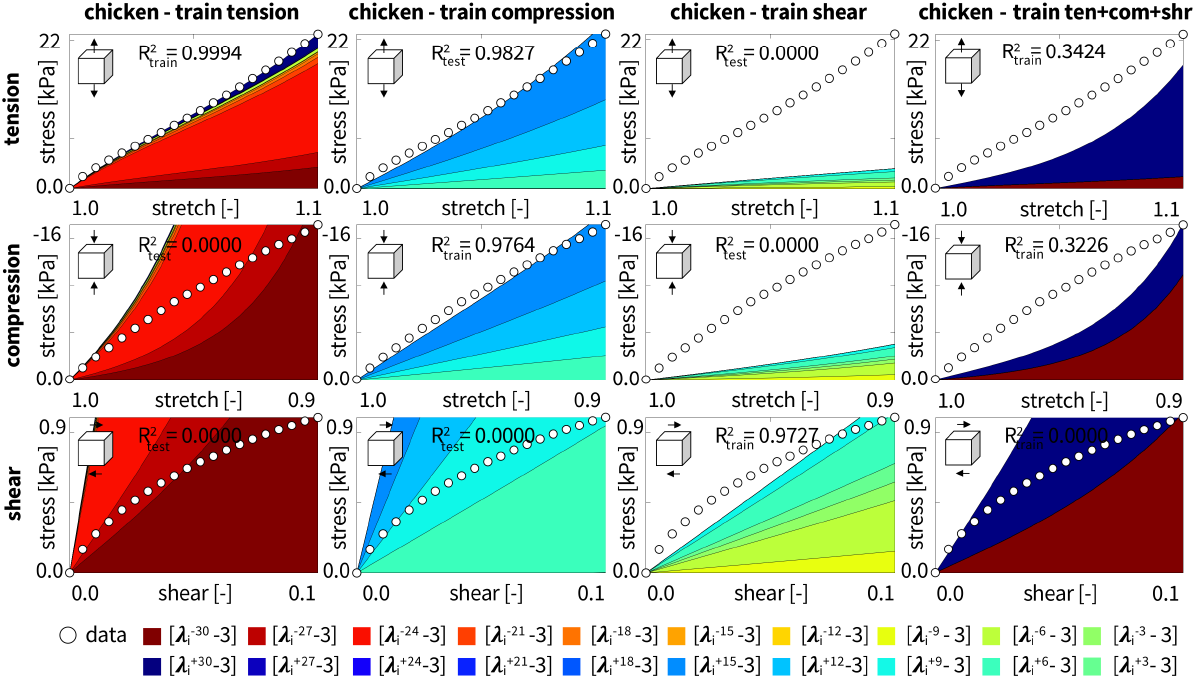
Real chicken data and and Ogden type model. Nominal stress as a function of stretch or shear strain for the network with 20 nodes from Figure 2. Training individually with tension, compression, or shear data from the real chicken, as well as with all three load cases simultaneously. Circles represent the experimental data. Color-coded regions designate the contributions of the 20 model terms to the stress function according to Figure 2. Coefficients of determination *R*^2^ indicate the goodness of fit for train and test data.

Second, for single-mode training, the model performs well in predicting the tension data for training on compression with 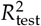 of 0.9827. Third, the network is unable to find a single model to fit the data for multi-mode training, with training fit 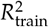of 0.3226, 0.3424, and 0.0000 for tension, compression, and shear. Fourth, for single-mode training with tension, the discovered terms are primarily negative red-type terms, for single-mode training with compression the terms are primarily positive blue-type terms, for single-mode training with shear there are both positive and negative terms, and multi-mode training also find both positive and negative terms. Lastly, all types of training discover a small subset of the twenty total possible terms even without additional regularization, with between two to seven terms visibly contributing to the stress function.

Multi-mode training discovers only two terms, the dark blue and dark red 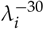 and 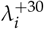 terms in the right column.

### 4.3 Valanis-Landel type mechanics of artificial and real meat

Figure 8 illustrates the automatically discovered constitutive models for tofurky using the fourteen term isotropic, perfectly incompressible, Valanis-Landel type network from Figure 3 with a penalty parameter of 0.001 for L2 regularization. First, for single-mode training, the model succeeds in fitting the individual sets of training data with 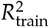values of 0.9993, 1.0000, and 0.9989 for tension, compression, and shear, respectively. In tension, the negative exponent trains to -14.42, while the positive exponent is not activated significantly. In compression, both exponents train to non-zero values, -11.43 and +13.14. In shear, they train to comparable values of -13.76 and +12.83. Second, for single-mode training, the model performs moderately well in predicting the tension data for training on compression with 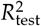 of 0.6908 and for predicting compression data for training on shear with 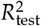 of 0.7257. Third, the network is able to find a good fit to the data for multi-mode training, with training fit 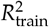of 0.8753, 0.9583, and 0.8268 for tension, compression, and shear, respectively. In multi-mode training, the exponents train to -23.25 and +16.43. Fourth, single mode training on tension data activates primarily the red-colored negative exponent while the other training modes activate both the red- and turquoise-colored exponents with large contributions to the stress function.

**Figure 8:**
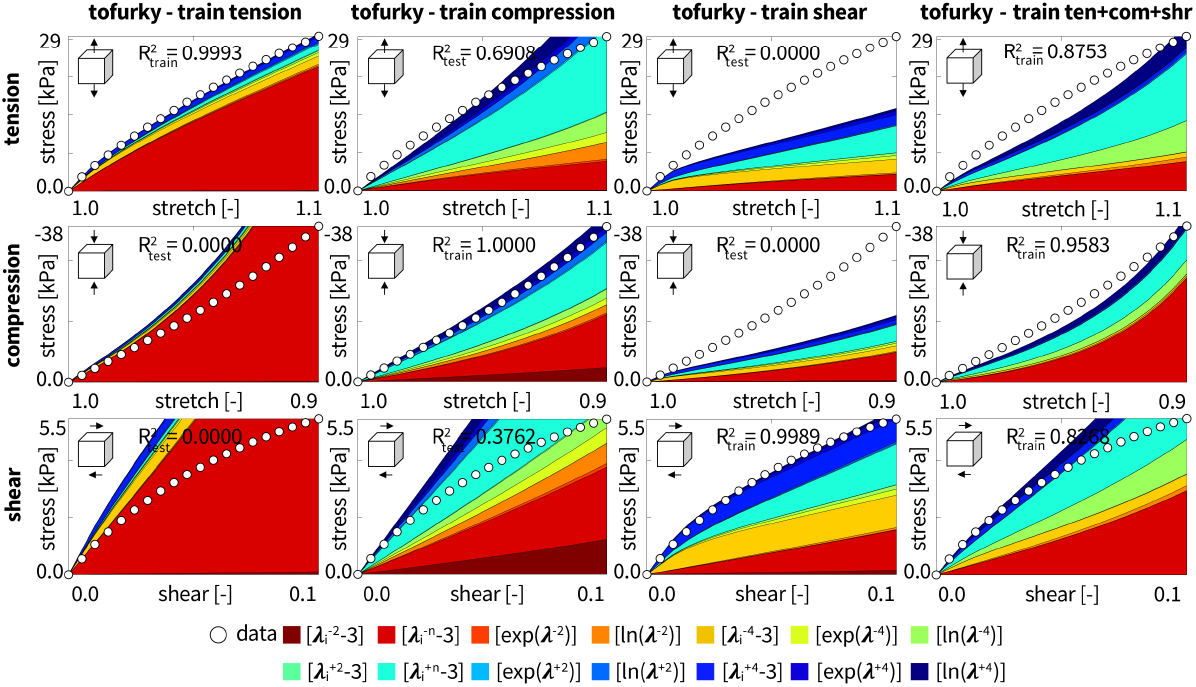
Tofurky data and Valanis-Landel type model. Nominal stress as a function of stretch or shear strain for the network with 14 nodes from Figure 3. Training individually with tension, compression, or shear data from the tofurky, as well as with all three load cases simultaneously, both with L2 regularization. Circles represent the experimental data. Color-coded regions designate the contributions of the 14 model terms to the stress function according to Figure 3. Coefficients of determination *R*^2^ indicate goodness of fit for train and test data.

Figure 9 illustrates the automatically discovered constitutive models for artificial chick’n using the fourteen term isotropic, perfectly incompressible, Valanis-Landel type network from Figure 3 with a penalty parameter of 0.001 for L2 regularization. Table 3 shows the discovered weights for multi-mode training. First, for single-mode training, the model succeeds in fitting the individual sets of training data with 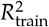values of 0.9998, 0.9867, and 0.9989 for tension, compression, and shear, respectively. In tension, the negative exponent trains to -12.89, while the positive exponent is not activated significantly. In compression, the positive exponent trains to +10.13, while the negative exponent is not activated significantly. In shear, both exponents contribute with comparable values of -9.23 and +9.19. Second, for single-mode training, the model performs moderately well in predicting the shear data for training on compression with 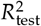of 0.8649 and for predicting compression data for training on shear with 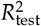 of 0.7257. Third, the network finds a moderately good fit to the data for multi-mode training, with training fit 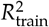of 0.5368, 0.9352, and 0.8345 for tension, compression, and shear, respectively. Fourth, multi-mode training does not activate either of the power terms and only discovers four terms that contribute to the stress function, the yellow 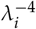 term, the green ln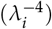 term, the blue 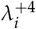 term, and the dark blue ln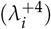 term in the right column.

**Table 3:**
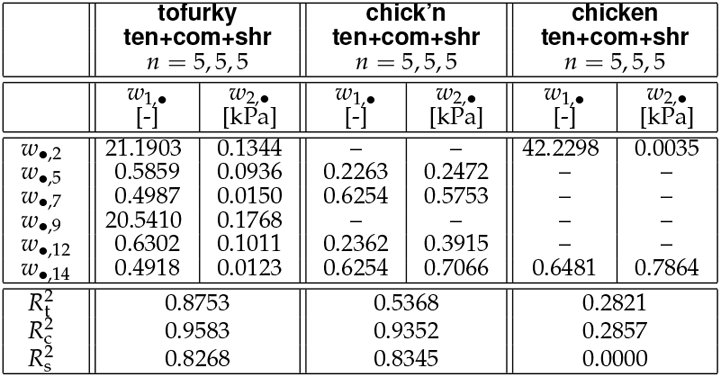
Tofurky, artificial chick’n, and real chicken and discovered Valanis-Landel type models and parameters. Models and parameters are discovered for simultaneous training with tension, compression, and shear data using the Valanis-Landel type neural network from Figure 3. Tofurky weights for regularization parameter *α* = 0.1 from Figure 11. Artificial chick’n and real chicken weights for *α* = 0.001 from Figures 9 and 10. Summary of the weights *w*_*k*_ and goodness of fit *R*^2^ for training in tension, compression, and shear.

| | tofurky<br>ten+com+shr<br>$n = 5, 5, 5$ | | chick'n<br>ten+com+shr<br>$n = 5, 5, 5$ | | chicken<br>ten+com+shr<br>$n = 5, 5, 5$ | |
| --- | --- | --- | --- | --- | --- | --- |
| | $w_{1,\bullet}$<br>[-] | $w_{2,\bullet}$<br>[kPa] | $w_{1,\bullet}$<br>[-] | $w_{2,\bullet}$<br>[kPa] | $w_{1,\bullet}$<br>[-] | $w_{2,\bullet}$<br>[kPa] |
| $w_{\bullet,2}$ | 21.1903 | 0.1344 | — | — | 42.2298 | 0.0035 |
| $w_{\bullet,5}$ | 0.5859 | 0.0936 | 0.2263 | 0.2472 | — | — |
| $w_{\bullet,7}$ | 0.4987 | 0.0150 | 0.6254 | 0.5753 | — | — |
| $w_{\bullet,9}$ | 20.5410 | 0.1768 | — | — | — | — |
| $w_{\bullet,12}$ | 0.6302 | 0.1011 | 0.2362 | 0.3915 | — | — |
| $w_{\bullet,14}$ | 0.4918 | 0.0123 | 0.6254 | 0.7066 | 0.6481 | 0.7864 |
| $R^2_t$ | 0.8753 | | 0.5368 | | 0.2821 | |
| $R^2_c$ | 0.9583 | | 0.9352 | | 0.2857 | |
| $R^2_s$ | 0.8268 | | 0.8345 | | 0.0000 | |

**Figure 9:**
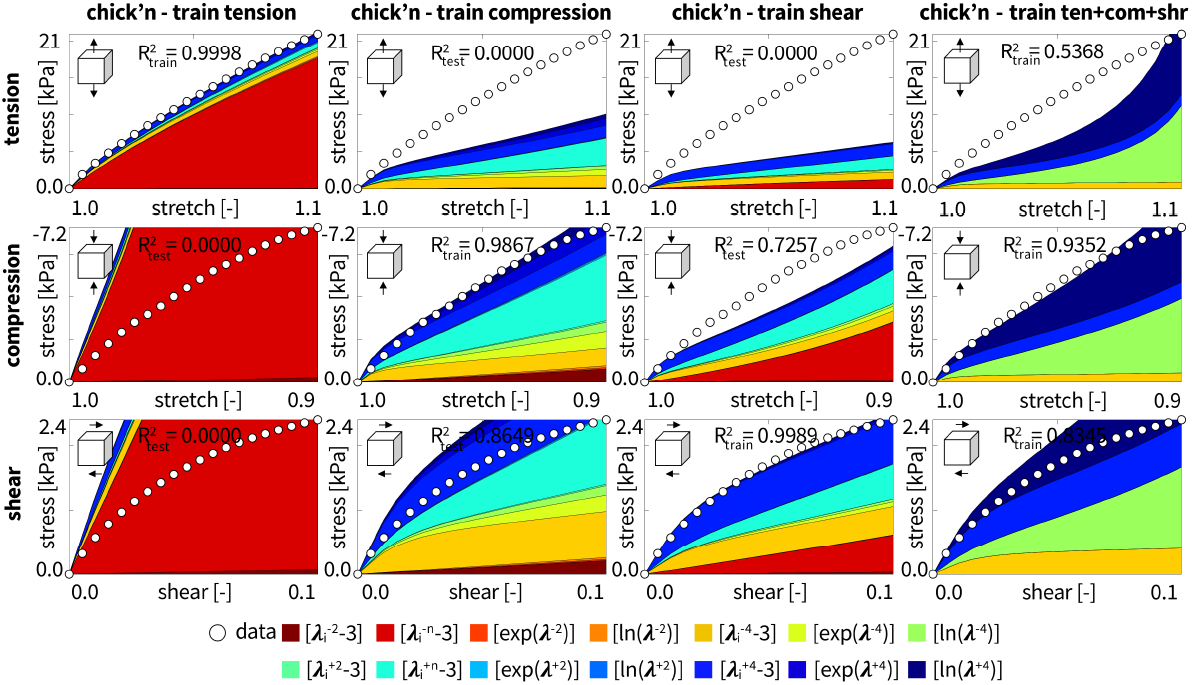
Artificial chick’n data and Valanis-Landel type model. Nominal stress as a function of stretch or shear strain for the network with 14 nodes from Figure 3. Training individually with tension, compression, or shear data from the artificial chick’n, as well as with all three load cases simultaneously, both with L2 regularization. Circles represent the experimental data. Color-coded regions designate the contributions of the 14 model terms to the stress function according to Figure 3. Coefficients of determination *R*^2^ indicate goodness of fit for train and test data.

Figure 10 illustrates the automatically discovered constitutive models for real chicken using the fourteen term isotropic, perfectly incompressible, Valanis-Landel type network from Figure 3 with a penalty parameter of 0.001 for L2 regularization. Table 3 shows the discovered weights for multi-mode training. First, for single-mode training, the model succeeds in fitting the individual sets of training data with 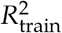values of 0.9999, 0.9949, and 0.9995 for tension, compression, and shear, respectively. In tension, the exponents train to -8.90 and +12.43. In compression, the positive exponent trains to +11.15, while the negative exponent is not activated significantly. In shear, both train to -8.32 and +9.96. Second, for single-mode training, the model performs well in predicting the compression data for training on tension with 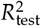of 0.9774 and in predicting tension data for training on compression with with 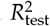of 0.9773. Third, the network is unable to find an adequate fit of the data for multi-mode training, with training fit 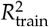 of 0.2821, 0.2857, and 0.0000 for tension, compression, and shear, respectively. In multi-mode training, the red trains to -42.23. Fourth, single-mode training discovers a wide range and number of terms while multimode training only discovers two terms, the red 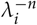 term and the dark blue ln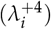term in the right column.

**Figure 10:**
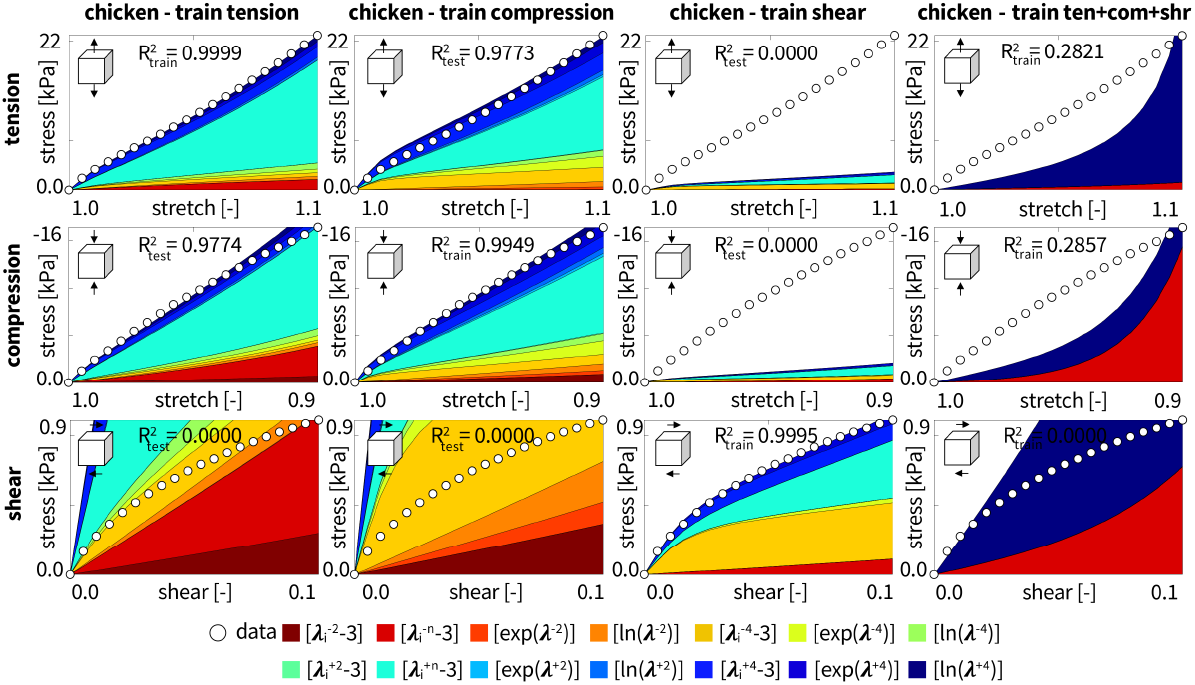
Real chicken data and Valanis-Landel type model. Nominal stress as a function of stretch or shear strain for the network with 14 nodes from Figure 3. Training individually with tension, compression, or shear data from the real chicken, as well as with all three load cases simultaneously, both with L2 regularization. Circles represent the experimental data. Color-coded regions designate the contributions of the 14 model terms to the stress function according to Figure 3. Coefficients of determination *R*^2^ indicate goodness of fit for train and test data.

Figure 11 illustrates the effect of added L2 regularization in multi-mode training for tofurky. The L2 penalty parameter *α* varies between 0.100, 0.010, 0.001, 0.000. Table 3 summarizes the discovered weights for multi-mode training with the *α* = 0.100 parameter. First, the variation in the number of discovered terms ranges from eight with zero penalty, on the right, to four with the largest penalty, on the left. With a penalty parameter of 0.1, the four discovered terms are the red 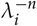 term, the turquoise 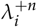 term, the yellow 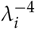 term, and the blue 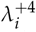 term, with the first two terms dominating in the left column. The exponents train to -21.19 and +20.54. Second, with an increasing penalty parameter *α*, the red and turquoise power terms contribute more to the stress function while all other terms contribute less. Third, increasing the penalty parameter has a negligible effect on the combined fit of the data with 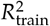values varying by less than 0.01 across all penalty parameters and loading modes.

**Figure 11:**
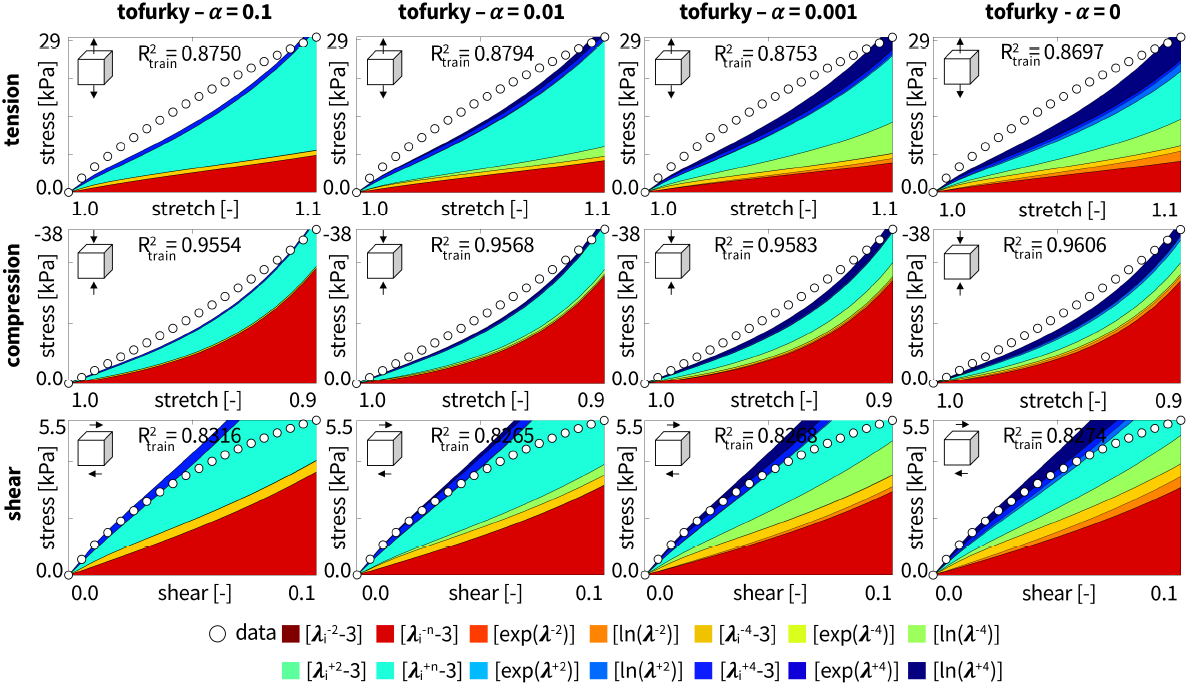
Tofurky data and Valanis-Landel type model with L2 regularization. Nominal stress as a function of stretch or shear strain for the network with 14 nodes from Figure 3. Training simultaneously with tension, compression, and shear data from the tofurky with L2 regularization with varying penalty parameters of 0.100, 0.010, 0.001, 0.000. Circles represent the experimental data. Color-coded regions designate the contributions of the 14 model terms to the stress function according to Figure 3. Coefficients of determination *R*^2^ indicate goodness of fit for train and test data.

### 4.4 Neo Hooke, Blatz Ko, and Mooney Rivlin mechanics of artificial and real meat

Figure 12 illustrates the automatically discovered parameters for tofurky when the Valanis-Landel type network from Figure 3 is restricted to the green neo Hooke term, 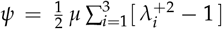 dark red Blatz Ko term, 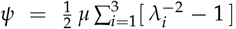 dark red and green Mooney Rivlin terms, 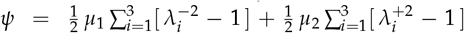 and red and turquoise Ogden terms, 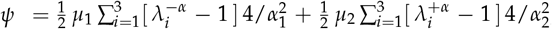 and simultaneously trained with tension, compression, and shear data. First, the neo Hooke, Blatz Ko, and Mooney Rivlin models all provide notably similar fits to the data, with 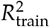values of 0.8777 or 0.8778, 0.9286, and 0.7329 or 0.7330 in tension, compression, and shear. Second, the two term Ogden model is able to discover a better simultaneous fit with a significant improvement for the shear data. The 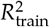values are 0.8717, 0.9734, and 0.8147. The negative Ogden exponent trains to *− α*_1_ = *−* 17.27, while the positive Ogden exponent trains to +*α*_2_ = +16.76.

**Figure 12:**
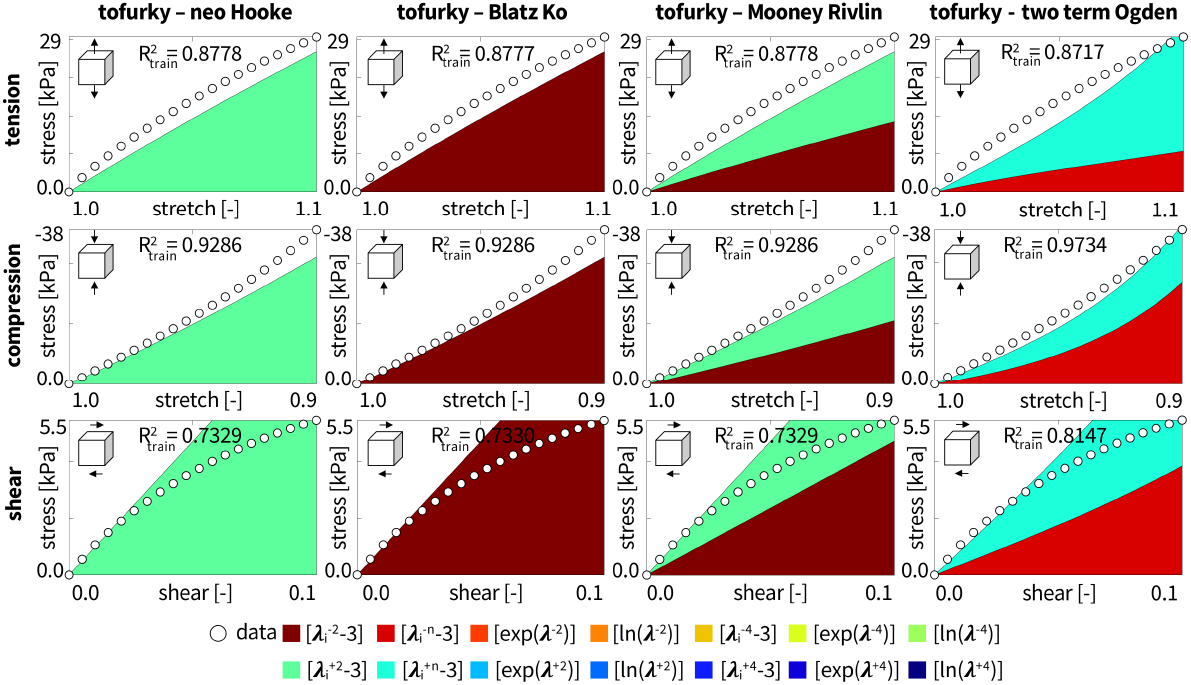
Tofurky data and special cases of neo Hooke, Blatz Ko, Mooney Rivlin, and two term Ogden models. Nominal stress as a function of stretch or shear strain for the Valanis-Landel type network from Figure 3 trained with all three load cases simultaneously. Circles represent the experimental data. The neo Hooke model uses only the +2 term, the Blatz Ko the -2 term, the Mooney Rivlin the -2 and +2 terms, and the two term Ogden the -n and +n terms. Color-coded regions designate the +2, -2, +2/-2, and +n/-n model terms to the stress function according to Fig. 3 multiplied by the weights from Table 4. Coefficients of determination *R*^2^ indicate goodness of fit for train data.

Table 4 shows the weights, shear moduli *µ*, and goodness of fit *R*^2^ when the Valanis-Landel type network from Figure 3 is restricted to the neo Hooke, Blatz Ko, Mooney Rivlin, or two *±n* Ogden terms and trained on all three loading modes simultaneously. The shear modulus of artificial chick’n is 35.6814 kPa, 35.6934 kPa, 35.6867 kPa, and 31.3620 kPa for the neo Hooke, Blatz Ko, Mooney Rivlin, and two term Ogden models. For comparison, the shear modulus of real chicken is 28.6748 kPa, 28.6731 kPa, 28.6709 kPa, and 14.3255 kPa for the neo Hooke, Blatz Ko, Mooney Rivlin, and two term Ogden models. First, while all models provide an adequate fit of the tension and compression tofurky data, the two term Ogden provides the best fit to the shear data with 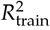 values of 0.8717, 0.9734, and 0.8147. Second, for artificial chick’n, the neo Hooke, Blatz Ko, and Mooney Rivlin models are not able to provide an adequate fit with 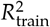 values of 0.0000, 0.0757 -0.0776, and 0.9124 -0.9126 for tension, compression, and shear. The two term Ogden models finds a better fit with 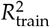 values of 0.6175, 0.7540, and 0.8501 in tension, compression, and shear. Third, none of the four models provide an adequate fit for real chicken. Of all four models, the two term Ogden provides the best fit with 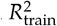 of 0.4013, 0.1676, and 0.0000 in tension, compression, and shear. Taken together, the two term Ogden model outperforms the neo Hooke, Blatz Ko, and Mooney Rivlin models for tofurky, artificial chick’n, and real chicken.

**Table 4:**
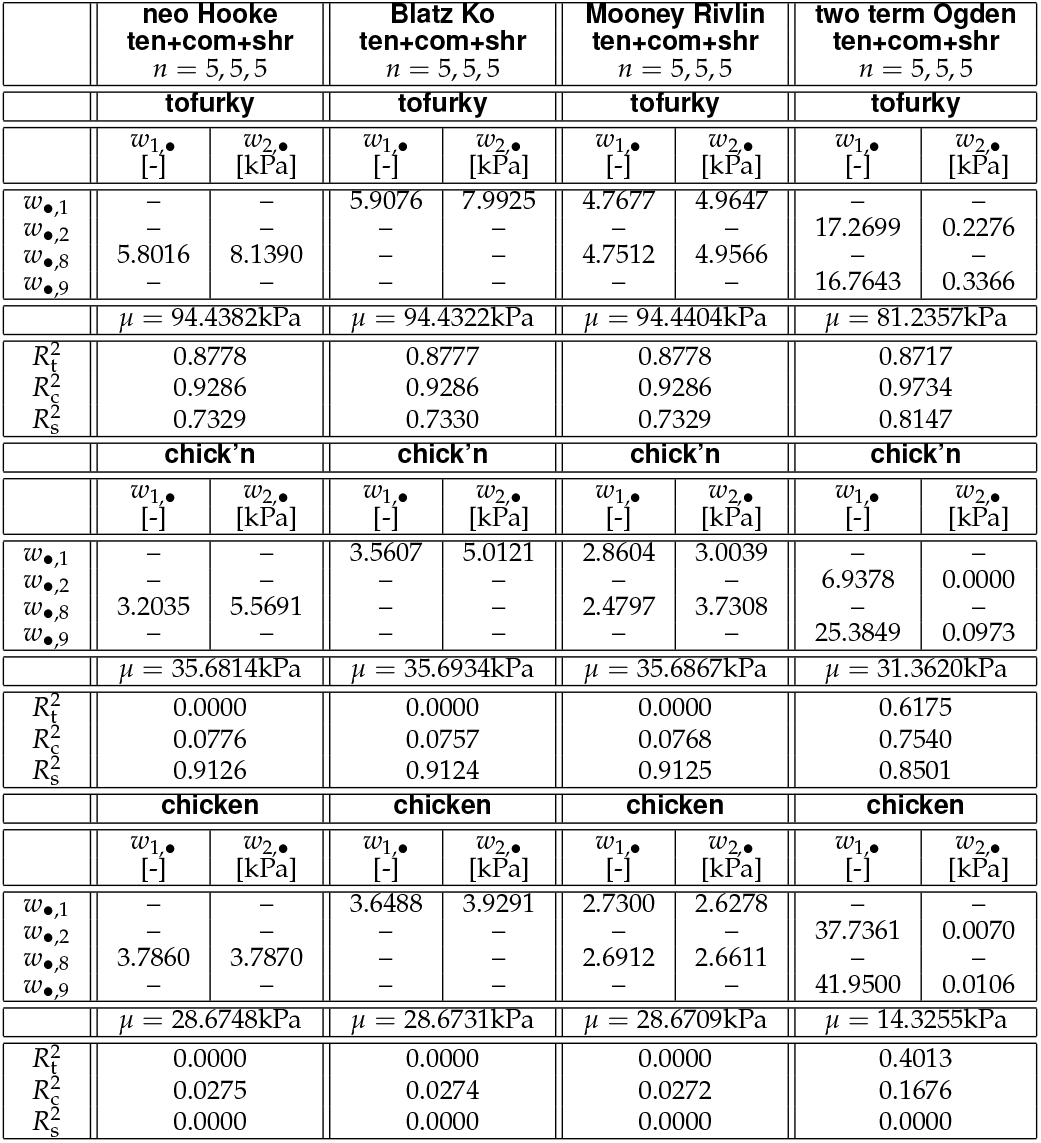
Special cases of neo Hooke, Blatz Ko, Mooney Rivlin and two term Ogden models. Tofurky, artificial chick’n, and real chicken parameters discovered for simultaneous training with tension, compression, and shear data. Summary of the non-zero weights from Figure 3 and detailed in equation (16), the shear moduli *µ*, and the goodness of fit *R*^2^ for the training data.

| | neo Hooke<br>ten+com+shr<br>$n = 5, 5, 5$ | | Blatz Ko<br>ten+com+shr<br>$n = 5, 5, 5$ | | Mooney Rivlin<br>ten+com+shr<br>$n = 5, 5, 5$ | | two term Ogden<br>ten+com+shr<br>$n = 5, 5, 5$ | |
| --- | --- | --- | --- | --- | --- | --- | --- | --- |
|  | tofurky |  | tofurky |  | tofurky |  | tofurky |  |
| | $w_{1,\bullet}$<br>[-] | $w_{2,\bullet}$<br>[kPa] | $w_{1,\bullet}$<br>[-] | $w_{2,\bullet}$<br>[kPa] | $w_{1,\bullet}$<br>[-] | $w_{2,\bullet}$<br>[kPa] | $w_{1,\bullet}$<br>[-] | $w_{2,\bullet}$<br>[kPa] |
| $w_{\bullet,1}$ | — | — | 5.9076 | 7.9925 | 4.7677 | 4.9647 | — | — |
| $w_{\bullet,2}$ | — | — | — | — | — | — | 17.2699 | 0.2276 |
| $w_{\bullet,8}$ | 5.8016 | 8.1390 | — | — | 4.7512 | 4.9566 | — | — |
| $w_{\bullet,9}$ | — | — | — | — | — | — | 16.7643 | 0.3366 |
| | $\mu = 94.4382\text{kPa}$ | | $\mu = 94.4322\text{kPa}$ | | $\mu = 94.4404\text{kPa}$ | | $\mu = 81.2357\text{kPa}$ | |
| $R_t^2$ | 0.8778 | | 0.8777 | | 0.8778 | | 0.8717 | |
| $R_c^2$ | 0.9286 | | 0.9286 | | 0.9286 | | 0.9734 | |
| $R_s^2$ | 0.7329 | | 0.7330 | | 0.7329 | | 0.8147 | |
|  | chick’n |  | chick’n |  | chick’n |  | chick’n |  |
| | $w_{1,\bullet}$<br>[-] | $w_{2,\bullet}$<br>[kPa] | $w_{1,\bullet}$<br>[-] | $w_{2,\bullet}$<br>[kPa] | $w_{1,\bullet}$<br>[-] | $w_{2,\bullet}$<br>[kPa] | $w_{1,\bullet}$<br>[-] | $w_{2,\bullet}$<br>[kPa] |
| $w_{\bullet,1}$ | — | — | 3.5607 | 5.0121 | 2.8604 | 3.0039 | — | — |
| $w_{\bullet,2}$ | — | — | — | — | — | — | 6.9378 | 0.0000 |
| $w_{\bullet,8}$ | 3.2035 | 5.5691 | — | — | 2.4797 | 3.7308 | — | — |
| $w_{\bullet,9}$ | — | — | — | — | — | — | 25.3849 | 0.0973 |
| | $\mu = 35.6814\text{kPa}$ | | $\mu = 35.6934\text{kPa}$ | | $\mu = 35.6867\text{kPa}$ | | $\mu = 31.3620\text{kPa}$ | |
| $R_t^2$ | 0.0000 | | 0.0000 | | 0.0000 | | 0.6175 | |
| $R_c^2$ | 0.0776 | | 0.0757 | | 0.0768 | | 0.7540 | |
| $R_s^2$ | 0.9126 | | 0.9124 | | 0.9125 | | 0.8501 | |
|  | chicken |  | chicken |  | chicken |  | chicken |  |
| | $w_{1,\bullet}$<br>[-] | $w_{2,\bullet}$<br>[kPa] | $w_{1,\bullet}$<br>[-] | $w_{2,\bullet}$<br>[kPa] | $w_{1,\bullet}$<br>[-] | $w_{2,\bullet}$<br>[kPa] | $w_{1,\bullet}$<br>[-] | $w_{2,\bullet}$<br>[kPa] |
| $w_{\bullet,1}$ | — | — | 3.6488 | 3.9291 | 2.7300 | 2.6278 | — | — |
| $w_{\bullet,2}$ | — | — | — | — | — | — | 37.7361 | 0.0070 |
| $w_{\bullet,8}$ | 3.7860 | 3.7870 | — | — | 2.6912 | 2.6611 | — | — |
| $w_{\bullet,9}$ | — | — | — | — | — | — | 41.9500 | 0.0106 |
| | $\mu = 28.6748\text{kPa}$ | | $\mu = 28.6731\text{kPa}$ | | $\mu = 28.6709\text{kPa}$ | | $\mu = 14.3255\text{kPa}$ | |
| $R_t^2$ | 0.0000 | | 0.0000 | | 0.0000 | | 0.4013 | |
| $R_c^2$ | 0.0275 | | 0.0274 | | 0.0272 | | 0.1676 | |
| $R_s^2$ | 0.0000 | | 0.0000 | | 0.0000 | | 0.0000 | |

## 5 Discussion

Artificial meat is marketed as an eco-friendly, plant-based protein that is an alternative to traditional meat sources but with similar taste and texture [15, 34]. Without a doubt, the mechanical properties of both real and artificial meat influence our eating experience [10, 36]. To quantify the mechanical properties of artificial meat, we investigated Tofurky® deli slices and Daring™ artificial chick’n, and compared them to real chicken. The tofurky was quite thin, only 1-1.5 mm thick on average, and was a rather isotropic, homogeneous material without distinct fiber directions. The artificial chick’n had distinct sheets, and we tested it in tension, compression, and shear parallel to the fiber direction as shown in Figure 1. For each meat type, we performed five tests per sample in tension, compression, and shear, and found that the standard error of the mean for each testing mode was small. This suggests that the mean of the tension, compression, and shear data is a robust characteristic of the material properties of each product.

### Constitutive neural networks automate the process of model discovery

To date, no unified constitutive model exists to characterize the mechanical behavior of artificial meat products. To discover a mathematical model that best describes the behavior of artificial and real meat in tension, compression, and shear, we compared two different neural networks, an Ogden type network, Figure 2, which we have previously successfully used to discover the best model for human brain tissue [40], and a new Valanis-Landel type network that includes some of the Ogden type features, but also exponential and logarithmic terms, Figure 3. Both networks performed well in single-mode fitting, and the Valanis-Landel model slightly outperformed the Ogden type model across all loading modes and all meat products. Both networks performed decently at predicting selective modes. The Ogden type network was able to predict shear from compression data for artificial chick’n and tension from compression data for real chicken. The Valanis-Landel type network was able to predict shear from compression data and compression from shear data for artificial chick’n and compression from tension data and tension from compression data for real chicken. For multi-mode training, the Ogden type network discovered a better overall fit to the data for real chicken, while the Valanis-Landel type network discovered a better model for artificial chick’n. The models performed equally well in simultaneously fitting the tofurky tension, compression, and shear data with a difference in cumulative 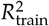values of less than 0.0001.

### Discovered models feature similar terms

The Ogden network features 20 different terms and discovers the best model out of a selection of 2^20^ *−*1 = 1, 048, 575 possible combinations of terms, meaning out of more than one million models. The Valanis Landel network features 14 different terms and discovers the best model out of a selection of 2^14^*−* 1 = 16, 383 possible combinations of terms, out of more than ten thousand models. Yet, when comparing both types of networks, we found that the Valanis Landel type network is generally more versatile: It contains a strong subset of Ogden type terms and spans a richer functional base overall. Interestingly, the Valanis Landel type network consistently discovers a similar subset of six terms for both artificial and real meat, while the weights of the other terms train to zero. All terms in the powers of two, 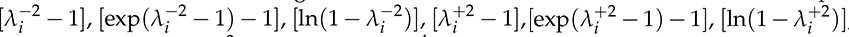, and all exponential terms, 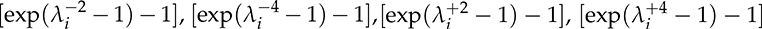, train to zero and are not present in any of the discovered models for combined tension, compression, and shear training. For tofurky, we robustly discover the general two term Ogden model with with flexible exponents in red and turquoise,

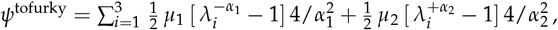

with two unit-less negative and positive exponents, *−α*_1_ = *−*21.19 and +*α*_2_ = +20.54, and two stiffness-like parameters, *µ*_1_ = 30.17 kPa and *µ*_2_ = 37.30 kPa. For artificial chick’n, we discover a four-term model in terms of the stretches to the negative and positive fourth power, 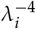 and 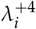 either plain, in yellow and blue, or combined with the classical Valanis Landel logarithm, in green and dark blue,

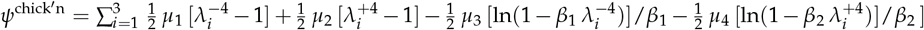

with two unit-less exponents *β*_1_ = 0.63 and *β*_2_ = 0.63 and four stiffness-like parameters *µ*_1_ = 0.11 kPa, *µ*_2_ = 0.18 kPa *µ*_3_ = 0.72 kPa, and *µ*_4_ = 0.88 kPa. For real chicken, we discover a two-terms model with one Ogden term with a negative exponent in red, and one Valanis Landel logarithm in dark blue,

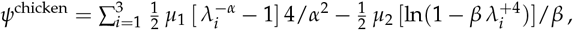

with two unit-less exponents *−α* = *−*42.23 and *β* = 0.65 and two stiffness-like parameters *µ*_1_ = 3.12 kPa and *µ*_2_ = 1.02 kPa. With the naked eye, and by touch, we immediately notice that artificial chick’n and real chicken are transversely isotropic with a single pronounced fiber direction. This creates an extremely low resistance to shear, about ten times lower than to axial loading which both our hyperelastic isotropic networks had difficulties to train on. In retrospect, it seems intuitive that we never discover any of the exponential terms: The concave shear softening nature of all three types of meat is in stark contrast to the convex strain stiffening behavior of collagenous soft tissues [6, 9, 19, 30], and rules out the exponential light red, light green, light blue, and dark blue terms of the Valanis Landel network for real and artificial meat. Strain softening seems to be much better represented by the red and turquoise classical Ogden-type terms with relatively large exponentials. At the same time, extreme exponentials on the order of *±*20 and above will likely not generalize well to stretch and shear ranges beyond the training regime of 10% deformation, where the stress contributions of these terms will simply explode. This implies that we need to be extremely careful when extrapolating beyond the training regime, especially when using Ogden type terms with large exponentials.

### Tofurky is three times stiffer than artificial chick’n and real chicken

When restricting the Valanis-Landel type network to its neo Hooke, Blatz Ko, Mooney Rivlin, and two-term Ogden model building blocks, and training on the tension, compression, and shear data simultaneously, we recover the classical shear modulus of tofurky, artificial chick’n, and real chicken. The neo Hooke, Blatz Ko, and Mooney Rivlin models all discovered identical shear moduli for tofurky with 94.4 kPa, artificial chick’n with 35.7 kPa, and real chicken with 21.4 kPa. Notably, we consistently discovered lower values for the two-term Ogden with shear moduli of tofurky with 81.2 kPa, artificial chick’n with 31.4 kPa, and real chicken with 14.3 kPa. This highlights that the choice of model has a significant impact on the mechanical moduli, even on the simplest of all, the shear modulus, which was up to 50% smaller when fit with the two-term Ogden model. Yet, the general trend remains the same across all four models: Of all three meat types, tofurky is the most isotropic and homogeneous and also displays the stiffest behavior. Both artificial chick’n and real chicken are anisotropic with a pronounced fiber direction, along which they display a similar behavior in tension, compression, and shear, and are only a third as stiff as tofurky.

### Limitations and future work

While we solidly and robustly discovered models and parameters for artificial and real meat products, our pilot study has a few limitations: First, we performed all tests on uncooked samples, but note that tofurky and artificial chick’n simply need to be warmed up whereas real chicken needs to be cooked. During cooking, the muscle fibers of real meat contract, cause a loss of water, and change the perceived tenderness of the meat [21]. Second, the tofurky deli slices are generally isotropic and homogeneous, but very thin. As a result, we tested the deli slices in the in-plane direction for tension and perpendicular to the in-plane direction for compression and shear, which might explain the varying stiffnesses for the different modes. Third, the artificial chick’n displays significant thickness variations and heterogeneous fiber sheets. When compressing the chick’n samples beyond 10%, the sheets collapsed on each other and the material response is no longer hyperelastic [22]. Fourth, the real chicken had a distinct fiber direction [46] that made it simple to cut similarly sized samples, but difficult to test in compression where chicken displayed a very soft, almost Jello-like consistency. Overall, to capture the anisotropy of artificial chick’n and real chicken, we would like to extend our network to include the fourth and fifth invariants [13, 31], similar to a previous network for skin [25]. Ideally, in the future, we would like to perform tension, compression, and shear experiments on the same samples using a single triaxial testing device, rather than using separate samples and separate instruments for separate testing modes [4]. We envision that performing additional tests will more robustly inform our network architecture, and allow us to combine successful terms of the Ogden type and Valanis-Landel type networks to a single set of terms and a single unified constitutive neural network to reliably and accurately discover the best model and parameters for artificial and real meat.

## 6 Conclusion

Plant-based meat substitutes are an increasingly popular alternative to animal consumption. The mechanical behavior of artificial meat directly influences our perception of taste through texture, consistency, and stiffness, and ultimately shapes our sensory experience and enjoyment of the product. However the mechanical properties of artificial meat and their comparison to real meat remain incompletely understood. Here we performed mechanical tension, compression, and shear tests on isotropic and anisotropic artificial and real meat and analyzed the data using custom-designed constitutive neural networks. Constitutive neural networks are a new, powerful and easy-to-use technology to discover the model and parameters that best explain a wide variety of soft matter systems. Especially for newly engineered materials like artificial meet for which the best model is not yet known, they allow us to rapidly screen thousands of potential model candidates and–entirely without bias and human interaction–select the best model. We compared two different constitutive neural networks, of Ogden type and Valanis-Landel type, and demonstrated that both can robustly fit the experimental data from tofurky, artificial chick’n, and real chicken in tension, compression, and shear. By design, both networks contain the classical neo Hooke, Blatz Ko, and Mooney Rivlin models as special cases for which they robustly discover shear moduli of 94.4 kPa for tofurky, 35.7 kPa for artificial chick’n, and 28.7 kPa for real chicken. This suggests that artificial chicken successfully reproduces the mechanical properties of real chicken across all loading modes, while tofurky does not and is about three times as stiff. We observed that all three meat products displayed shear softening with concave stress-stretch curves, and, more importantly, that their resistance to shear was about an order of magnitude lower than their resistance to tension and compression. Our new neural network-based automated model discovery can quantify the mechanics of artificial and real meat and has the potential to inform the design of plant-based meal substitutes. Understanding the mechanics of artificial meat has the potential to revolutionize food production, address environmental concerns, and ensure a sustainable future for our planet.

## Data Availability

Our source code, data, and examples are available at https://github.com/LivingMatterLab/CANN.

## Acknowledgments

This work was supported by a NSF Graduate Research Fellowship to Skyler St. Pierre, and by the Stanford School of Engineering Covid-19 Research and Assistance Fund and Stanford Bio-X IIP seed grant to Ellen Kuhl.

## References

[1] Alber, M., Buganza Tepole, A., Cannon, W., De, S., Dura-Bernal, S., Garikipati, K., Karniadakis, G.E., Lytton, W.W., Perdikaris, P., Petzold, L., Kuhl, E., 2019. Integrating machine learning and multiscale modeling: Perspectives, challenges, and opportunities in the biological, biomedical, and behavioral sciences. npj Digital Medicine 2, 115. doi:10.1038/s41746-019-0193-y.

[2] As’ad, F., Avery, P., Farhat, C., 2022. A mechanics-informed artificial neural network approach in data-driven constitutive modeling. International Journal for Numerical Methods in Engineering 123, 2738–2759. doi:10.1002/nme.6957.

[3] Blatz, P.J., Ko, W.L., 1962. Application of finite elastic theory to the deformation of rubbery materials. Transactions of the Society of Rheology 6, 223–252. doi:10.1122/1.548937.

[4] Budday, S., Sommer, G., Birkl, C., Langkammer, C., Haybaeck, J., Kohnert, J., Bauer, M., Paulsen, F., Steinmann, P., Kuhl, E., Holzapfel, G.A., 2017. Mechanical characterization of human brain tissue. Acta Biomaterialia 48, 319–340. doi:10.1016/j.actbio.2016.10.036.

[5] Clifford Astbury, C., 2023. Health and sustainability of everyday food. Nature Food 4, 357. doi:10.1038/s43016-023-00761-6.

[6] De Kegel, D., Vastmans, J., Fehervary, H., Depreitere, B., Vander Sloten, J., Famaey, N., 2018. Biomechanical characterization of human dura mater. Journal of the Mechanical Behavior of Biomedical Materials 79, 122–134. doi:10.1016/j.jmbbm.2017.12.023.

[7] Dekkers, B.L., Boom, R.M., van der Goot, A.J., 2018. Structuring processes for meat analogues. Trends in Food Science & Technology 81, 25–36. doi:10.1016/j.tifs.2018.08.011.

[8] Eggersmann, R., Stainier, L., Ortiz, M., Reese, S., 2021. Model-free data-driven computational mechanics enhanced by tensor voting. Computer Methods in Applied Mechanics and Engineering 373, 113499. doi:10.1016/j.cma.2020.113499.

[9] Famaey, N., Sommer, G., Vander Sloten, J., Holzapfel, G.A., 2012. Arterial clamping: finite element simulation and in vivo validation. Journal of the Mechanical Behavior of Biomedical Materials 12, 107–118. doi:10.1016/j.jmbbm.2012.03.010.

[10] Fiorentini, M., Kinchla, A.J., Nolden, A.A., 2020. Role of sensory evaluation in consumer acceptance of plant-based meat analogs and meat extenders: A scoping review. Foods 9. doi:10.3390/foods9091334.

[11] Flaschel, M., Kumar, S., De Lorenzis, L., 2021. Unsupervised discovery of interpretable hyperelastic constitutive laws. Computer Methods in Applied Mechanics and Engineering 381, 113852.

[12] Flaschel, M., Yu, H., Reiter, N., Hinrichsen, J., Budday, S., Steinmann, P., Kumar, S., De Lorenzis, L., 2023. Automated discovery of interpretable hyperelastic material models for human brain tissue with EUCLID. arXiv doi:10.48550/arXiv.2305.16362.

[13] Fuhg, J.N., Bouklas, N., 2022. On physics-informed data-driven isotropic and anisotropic constitutive models through probabilistic machine learning and space-filling sampling. Computer Methods in Applied Mechanics and Engineering 394, 114915.

[14] Fuhg, J.N., Bouklas, N., Jones, R.E., 2022. Learning hyperelastic anisotropy from data via a tensor basis neural network. Journal of the Mechanics and Physics of Solids 168, 105022.

[15] Hartmann, C., Siegrist, M., 2017. Consumer perception and behaviour regarding sustainable protein consumption: A systematic review. Trends in Food Science & Technology 61, 11–25. doi:10.1016/j.tifs.2016.12.006.

[16] Hartmann, S., 2001. Parameter estimation of hyperelasticity relations of generalized polynomial-type with constraint conditions. International Journal of Solids and Structures 38, 7999–8018.

[17] He, J., Evans, N.M., Liu, H., Shao, S., 2020. A review of research on plant-based meat alternatives: Driving forces, history, manufacturing, and consumer attitudes. Comprehensive Reviews in Food Science and Food Safety 19, 2639–2656. doi:10.1111/1541-4337.12610.

[18] Hinrichsen, J., Reiter, N., Bräuer, L., Paulsen, F., Kaessmair, S., Budday, S., 2022. Inverse identification of region-specific hyperelastic material parameters for human brain tissue. bioRxiv doi:10.1101/2022.12.19.521022.

[19] Holzapfel, G., 2000. Nonlinear Solid Mechanics: A Continuum Approach for Engineering. Wiley. doi:10.1023/A:1020843529530.

[20] Holzapfel, G.A., Linka, K., Sherifova, S., Cyron, C., 2021. Predictive constitutive modelling of arteries by deep learning. Journal of the Royal Society Interface 18, 20210411. doi:10.1098/rsif.2021.0411.

[21] Ježek, F., Kameník, J., Macharáčková, B., Bogdanovičová, K., Bednář, J., et al., 2020. Cooking of meat: effect on texture, cooking loss and microbiological quality–a review. Acta Veterinaria Brno 88, 487–496. doi:10.2754/avb201988040487.

[22] Jonkers, N., van Dijk, W.J., Vonk, N.H., van Dommelen, J.A.W., Geers, M.G.D., 2022a. Anisotropic mechanical properties of Selective Laser Sintered starch-based food. Journal of Food Engineering 318, 110890. doi:10.1016/j.jfoodeng.2021.110890.

[23] Jonkers, N., van Dommelen, J.A.W., Geers, M.G.D., 2022b. Intrinsic mechanical properties of food in relation to texture parameters. Mechanics of Time-Dependent Materials 26, 323–346. doi:10.1007/s11043-021-09490-4.

[24] Klein, D.K., Fernandez, M., Martin, R.J., Neff, P., Weeger, O., 2022. Polyconvex anisotropic hyperelasticity with neural networks. Journal of the Mechanics and Physics of Solids 159, 105703.

[25] Linka, K., Buganza Tepole, A., Holzapfel, G.A., Kuhl, E., 2023a. Automated model discovery for skin: Discovering the best model, data, and experiment. Computer Methods in Applied Mechanics and Engineering 410, 116007. doi:10.1101/2022.12.19.520979.

[26] Linka, K., Hillgärtner, M., Abdolazizi, K.P., Aydin, R.C., Itskov, M., Cyron, C.J., 2021. Constitutive artificial neural networks: A fast and general approach to predictive data-driven constitutive modeling by deep learning. Journal of Computational Physics 429, 110010. doi:10.1016/j.jcp.2020.110010.

[27] Linka, K., Kuhl, E., 2023. A new family of Constitutive Artificial Neural Networks towards automated model discovery. Computer Methods in Applied Mechanics and Engineering 403, 115731. doi:10.1016/j.cma.2022.115731.

[28] Linka, K., St. Pierre, S.R., Kuhl, E., 2023b. Automated model discovery for human brain using constitutive artificial neural networks. Acta Biomaterialia 160, 134–1510. doi:10.1016/j.actbio.2023.01.055.

[29] Masi, F., Stefanou, I., Vannucci, P., Maffi-Berthier, V., 2021. Thermodynamics-based artificial neural networks for constitutive modeling. Journal of the Mechanics and Physics of Solids 147, 04277.

[30] Menzel, A., 2006. A fibre reorientation model for orthotropic multiplicative growth. Biomechanics and Modeling in Mechanobiology 6, 303–320. doi:10.1007/s10237-006-0061-y.

[31] Menzel, A., Steinmann, P., 2003. A view on anisotropic finite hyper-elasticity. European Journal of Mechanics - A/Solids 22, 71–87. doi:10.1016/S0997-7538(02)01253-6.

[32] Mooney, M., 1940. A theory of large elastic deformation. Journal of applied physics 11, 582–592. doi:10.1063/1.1712836.

[33] Mullen, A., 2022. The price is right for artificial meat. Nature Food 3, 813. doi:10.1038/s43016-022-00629-1.

[34] Nezlek, J.B., Forestell, C.A., 2022. Meat substitutes: Current status, potential benefits, and remaining challenges. Current Opinion in Food Science 47, 1008–1090. doi:10.1016/j.cofs.2022.100890.

[35] Ogden, R.W., 1972. Large deformation isotropic elasticity–on the correlation of theory and experiment for incompressible rubberlike solids. Proceedings of the Royal Society of London. A. Mathematical and Physical Sciences 326, 565–584. doi:10.1098/rspa.1972.0026.

[36] Pascua, Y., Koç, H., Foegeding, E.A., 2013. Food structure: Roles of mechanical properties and oral processing in determining sensory texture of soft materials. Current Opinion in Colloid & Interface Science 18, 324–333. doi:10.1016/j.cocis.2013.03.009.

[37] Post, M.J., Levenberg, S.L., Kaplan, D.L., Genovese, N.G., Fu, J., Bryant, C.J., Negowetti, N., Verzijden, K., Moutsatsous, P., 2020. Scientific, sustainability and regulatory challenges of cultured meat. Nature Food 1, 403–415. doi:10.1038/s43016-020-0112-z.

[38] Rezaei, S., Harandi, A., Moeineddin, A., Xu, B.X., S., R., 2022. A mixed formulation for physics-informed neural networks as a potential solver for engineering problems in heterogeneous domains: comparison with finite element method. Computer Methods in Applied Mechanics and Engineering 401, 115616. doi:10.1016/j.cma.2022.115616.

[39] Rivlin, R.S., 1948. Large elastic deformations of isotropic materials iv. further developments of the general theory. Philosophical transactions of the royal society of London. Series A, Mathematical and physical sciences 241, 379–397. doi:10.1098/rsta.1948.0024.

[40] St. Pierre, S.R., Linka, K., Kuhl, E., 2023. Principal-stretch-based constitutive neural networks autonomously discover a subclass of ogden models for human brain tissue. Brain Multiphysics 4, 100066. doi:10.1101/2023.01.14.524079.

[41] Tac, V., Sahli Costabal, F., Buganza Tepole, A., 2022. Data-driven tissue mechanics with polyconvex neural ordinary differential equations. Computer Methods in Applied Mechanics and Engineering 398, 115248. doi:10.1016/j.cma.2022.115248.

[42] Treloar, L., 1948. Stresses and birefringence in rubber subjected to general homogeneous strain. Proceedings of the Physical Society (1926-1948) 60, 135. doi:10.1088/0959-5309/60/2/303.

[43] UBS Global, 2019. The food revolution. URL: https://www.ubs.com/global/en/wealth-management/insights/chief-investment-office/sustainable-investing/2019/food-revolution.html.

[44] Valanis, K., Landel, R.F., 1967. The strain-energy function of a hyperelastic material in terms of the extension ratios. Journal of Applied Physics 38, 2997–3002. doi:10.1063/1.1710039.

[45] Valanis, K.C., 2022. The valanis–landel strain energy function elasticity of incompressible and compressible rubber-like materials. International Journal of Solids and Structures 238, 111271. doi:10.1016/j.ijsolstr.2021.111271.

[46] Wang, L.M., Linka, K., Kuhl, E., 2023. Automated model discovery for muscle using constitutive recurrent neural networks. bioRxiv doi:10.1101/2023.05.09.540027.

